# Acquisition, Evolution, and Diversification of Genomic Islands: A Case Study of a Virulence Gene Cluster in *Pseudomonas syringae*

**DOI:** 10.1101/2024.06.12.598671

**Authors:** Luiz Orlando de Oliveira, Hugo Vianna Silva Rody, Selene Aguilera, Jesús Murillo

## Abstract

Genomic islands are widely distributed among bacteria, facilitating the dissemination of genes relevant to human, animal, and plant health. Herein, we explore the evolutionary history of a genomic island (Tox island) composed of a novel mobile genetic element (GInt) carrying an ∼25 kb gene cluster (Pht cluster) responsible for the biosynthesis of the phytotoxin phaseolotoxin, a virulence factor of the plant pathogen *Pseudomonas syringae*. The Pht cluster has been acquired, either naked or associated with a GInt, on seven independent occasions by four phylogroups of *P. syringae* and the distant rhizobacterium *Pseudomonas* sp. JAI115. The Pht cluster was independently captured by three distinct GInt elements, suggesting specific mechanisms for gene capture. Once acquired, the Tox island tends to be stably maintained, evolving with the genome. An array of molecular analyses delineated the likely evolutionary trajectory of the Pht cluster within *P. syringae* pv. actinidiae (Psa) and *P. amygdali* pv. phaseolicola (Pph), involving: 1) acquisition by Pph; 2) transfer of haplotype G to Psa biovar 1; 3) acquisition or replacement by a haplotype of haplogroup D in Psa biovar 1; 4) acquisition of haplotype C by Psa biovar 6; and 5) replacement of the Tox island in Pph by a distantly related GInt. These findings underscore the potential role of phaseolotoxin in bacterial fitness and contribute to our understanding of virulence evolution in plant pathogens. Furthermore, GInts provide a model for studying the evolutionary dynamics of mobile genetic elements and the dissemination of adaptive genes among bacterial pathogens.

## Introduction

Genomic islands (GIs) are loosely defined as discrete DNA regions that are present only in certain individuals of a bacterial lineage and are likely acquired from other organisms (Darmon and Leach, 2014). GIs are usually large, from 10 to more than 250 kb, with their cargo DNA allowing the rapid evolution of bacteria and leading to their adaptation to stressful environments and occupation of new niches. GIs are particularly important in the spread of resistance to antibiotics and heavy metals, acquisition of pathogenicity, and biosynthesis of secondary metabolites.

GIs are highly diverse in origin, structure, and size and can either be “naked” GIs, comprising a stretch of DNA that is not associated with any obvious mobile gene, or result from the activity of different mobile genetic elements (MGEs) with distinct structures and associated DNA (Audrey et al., 2023). Examples of MGEs that appear as GIs include integrative and conjugative elements (ICEs), prophages, transposons, and integrated plasmids. Although ICEs and related elements have been extensively investigated (Bellanger et al., 2014), most GIs in bacteria are either naked or generated by other types of MGEs, nearly one-third of which are associated with tyrosine integrases (Audrey et al., 2023). However, many of these elements are at most poorly characterized, resulting in our inability to estimate or predict their potential impact on the evolution of bacteria and the horizontal dissemination of adaptive genes. Due to their sheer abundance (Audrey et al., 2023; Idola et al., 2023), all these elements constitute a vast and unexplored source of genetic variation in bacteria, whose activity can greatly impact human wellbeing.

GInts are a new class of integrative MGE that are defined by their distinct tripartite structure: 1) a four-gene operon (*ginABCD*) at their 5’ end, 2) a cargo DNA region that can be as large as 70 kb, and 3) a very short and poorly conserved 3’ end, although GInts are bordered by short repeats (*attL* and *attR*) generated during the integration process and typical of integrative elements (Bardaji et al., 2017). The *gin* operon encodes three tyrosine integrases and a putative helix-turn-helix protein, which are all essential for mobilization. GInts excise with low frequency as circular intermediate molecules, which are likely unable to replicate autonomously, often perfectly reconstructing the insertion site in the genome and leaving no scar. Phylogenetic reconstructions clearly showed that GInts are horizontally transferred among bacteria; however, they do not carry determinants for conjugative transfer and have not been shown to be associated with plasmids. Currently, transfer of the element has not been reproduced in the laboratory, and it is not clear how it is mobilized in the bacterial population.

GInts can display high insertion specificity, and the integration and excision processes may direct GInt evolution toward greater host specialization, with a preference for phylogenetically related hosts, as occurs with other MGEs (Jackson et al., 2011; Bardaji et al., 2017). There is low overall sequence conservation among the *gin* operons of different GInt families, which limits the ability to identify GInts solely based on sequence similarity (Bardaji et al., 2017; Idola et al., 2023). However, the presence of the *gin* operon is a distinctive and defining feature of GInts and can be used as a marker to identify these elements in bacterial genomes. Additionally, bacterial genomes can contain several dissimilar GInts inserted in different locations. The cargo DNA is also highly variable in gene content and size, is preferentially captured from phylogenetically related bacteria, and often confers specific functions that could provide an adaptive advantage to the bacterial host (Bardaji et al., 2017). Examples of cargo DNA include virulence genes, antibiotic resistance gene clusters, and gene clusters involved in complex phenotypes such as toxin biosynthesis. The so-called ‘Tox island’ of *Pseudomonas syringae* illustrates the complexity of biosynthetic pathways that GInts may deliver in their cargo DNA. In the Tox island, the cargo DNA consists of a large gene cluster involved in the full biosynthesis of phaseolotoxin (Genka et al., 2006; Aguilera et al., 2007; Bardaji et al., 2017).

*P. syringae* is a bacterial complex of species that causes plant diseases with significant economic impacts and is a relevant model in plant‒microbe research (Mansfield et al., 2012). The *P. syringae* complex includes strains previously classified into 15 different species and has been divided into 13 different phylogroups that can be classified as separate species (Berge et al., 2014; Gomila et al., 2017; Dillon et al., 2019). Additionally, *P. syringae* has been divided into pathovars depending on the plant host range and pathogenic characteristics. *P. syringae* strains can produce one or more phytotoxins, which are generally important components of their virulence arsenal (Bender et al., 1999).

Phaseolotoxin is a tripeptide toxin that inhibits ornithine carbamoyltransferase (Fig. S1), leading to a depletion of arginine and the death of target eukaryotic and prokaryotic cells (Bender et al., 1999). This toxin is responsible for the characteristic halos produced in infected plants (Mitchell, 1976; Tamura et al., 2002). Although definitive evidence is lacking, the available data suggest that phaseolotoxin contributes to the systemic infection of the plant host and the intensity of disease symptoms; therefore, the toxin is considered a virulence factor (Patil et al., 1974; Arndt et al., 1989; Miyoshi et al., 2012; Sawada and Fujikawa, 2019). Additionally, plant extracts stimulate toxin biosynthesis (Hernández-Morales et al., 2009), supporting its role in the pathogenic process. The production of phaseolotoxin has been demonstrated only for strains of the *P. syringae* complex, and belonging to *P. amygdali* pv. phaseolicola (Pph) and *P. syringae* pv. actinidiae biovars 1 (Psa1) and 6 (Psa6), as well as by *P. syringae* pv. syringae strain CFBP3388 (Mitchell, 1978; Tourte and Manceau, 1995; Tamura et al., 2002; Murillo et al., 2011).

Phaseolotoxin biosynthesis and resistance depend on a cluster of 23 genes, the Pht cluster, which is carried by a GInt (Genka et al., 2006; Aguilera et al., 2007; Bardaji et al., 2017). Remarkably, the Pht cluster is practically identical in strains of the phylogenetically distant Pph and Psa, which, together with phylogenetic analyses, indicates that the Pht cluster has been acquired by horizontal transfer (Sawada et al., 2002). However, previous studies have not clarified the evolutionary history of the Pht cluster within the *P. syringae* complex.

Despite the importance of phytotoxins, such as phaseolotoxin, in the pathogenic process (e.g., Sawada and Fujikawa, 2019; Uddin et al., 2022), their evolutionary histories are not well understood. Additionally, little is known about the routes of acquisition of biosynthetic genes and their dissemination within bacterial populations.

Here, we used molecular phylogeny, gene genealogy, and network analyses to uncover the complex history of MGE capture and horizontal transfer of the Pht cluster. We initiated our study by exploring the genealogical relationships and diversification levels among the extant Pht clusters within the *P. syringae* complex. We assessed whether the Pht cluster sequences represented a single lineage or whether multiple independent lineages better explained our findings. Subsequently, we examined the genomic context surrounding integration events to determine whether the integration of Pht clusters resulted from a single horizontal transfer event within an ancestral strain, followed by diversification. We anticipated that congruence between the phylogenies of the Pht cluster and the chromosomal locus housing it would suggest a single acquisition event. Alternatively, incongruence may indicate multiple independent acquisition events accounting for the occurrence of Pht clusters in distinct host lineages. Finally, we investigated whether a single lineage of GInts facilitated the integration of all extant Pht clusters into host genomes or whether distinct lineages of GInts were involved. This exploration allowed us to consider diverse mechanisms of Pht cluster acquisition and diversification within the *P. syringae* complex.

## Results

### Insertion sites of the Pht cluster in *P. syringae* strains

The genome of Psy strain CFBPP3388 contained a Pht cluster similar to those of Pph and Psa, with homology ending after the stop codons of *argK* and *phtV* (Fig. 1). The Pht cluster in CFBP3388 was not part of a GInt and was inserted in a different genomic region than that in Pph and Psa. Although an IS*5*-family insertion sequence followed *phtV*, we found no other repeated elements or tRNA near the Pht cluster.

**Figure 1.**
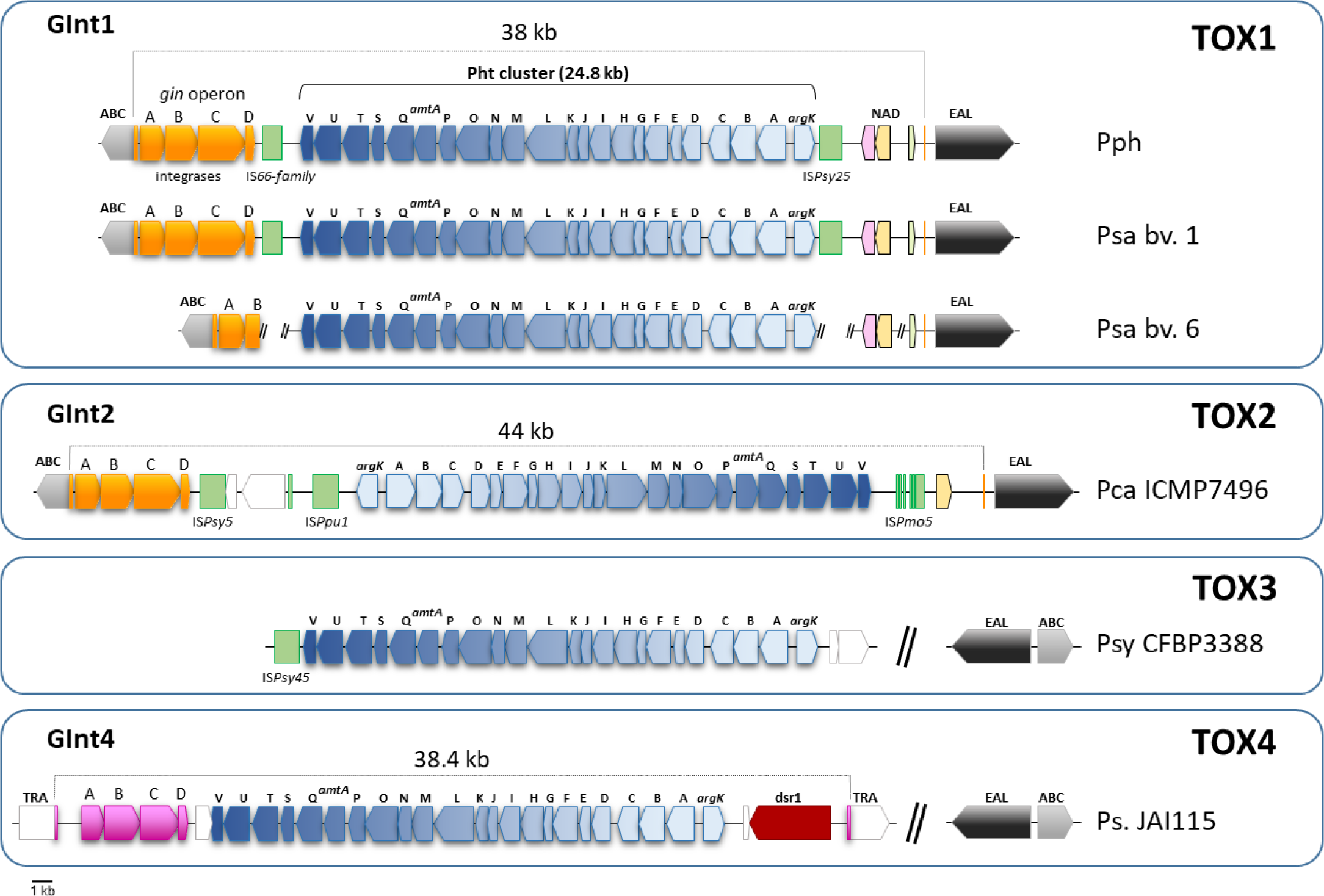
The four genomic configurations (TOX1 to TOX4) of the phaseolotoxin biosynthesis genes (Pht cluster) in toxigenic bacteria of the *Pseudomonas syringae* complex. Coding sequences, and their orientation, are indicated by block arrows, with identical colors indicating homology of sequence. Insertion sequences are indicated by green blocks, irrespective of homology. Overall orientation of the insertion of the Pht cluster is depicted in a blue spectrum for reference purposes. The gin operon (*ginABCD*) encodes three integrases and a putative helix-turn-helix protein that are essential for excision and integration of the transposon; IS, insertion sequence; ABC, putative ABC transporter (e.g., WP_011169501.1); NAD, NAD-dependent protein deacylase (e.g., WP_003381720.1); EAL, EAL domain-containing protein (e.g., WP_017698913.1); TRA, transporter substrate-binding domain-containing protein (WP_260319991.1); dsr1, anti-phage defense-associated sirtuin Dsr1 (WP_183785383.1). Abbreviations: Pph, *Pseudomonas amygdali* pv. phaseolicola; Psa, *P. syringae* pv. actinidiae; Pca, *P. caricapapayae*; Psy, *P. syringae* pv. syringae. All features are drawn to scale.

We also found a complete Pht cluster in the genome of *P. caricapapayae* (Pca) strain ICMP7496, which was carried by a GInt homologous to that of Pph and Pca and inserted in the same place: the 5’ end of an ABC transporter (Fig. 1). However, the DNA cargo of both GInts differed, and the Pht cluster in Pca was inverted compared to that in Pph and Psa (Fig. 1). An *E. coli* growth inhibition assay confirmed phaseolotoxin production by Pca CFBP4336 (syn. ICMP7496) at 18 °C and 28 °C (Fig. S2), confirming the functionality of the Pht cluster in this strain.

### The Pht cluster branches into three clades in *P. syringae*

We examined the presence of the 23 Pht cluster genes in 329 *P. syringae* strains, which represent six phylogroups and included all known phaseolotoxin-producing strains.

Our analysis identified 101 toxigenic strains with DNA homologous to the complete Pht cluster originally described in Pph and Psa (Genka et al., 2006; Aguilera et al., 2007). This included Pph, Psa1 and Psa6, Pca ICMP7496, and Psy CFBP3388 and 3023. Additionally, this included nine *P. syringae* pv. mellea and *P. syringae* pv. glycinea strains, as well as *Pseudomonas* sp. BDAL1 and KBS0707, all of which clustered with Pph in core genome phylogenies and are likely misclassified Pph strains. The complete Pht cluster spanned from PSPPH_4319 (*argK*) to PSPPH_4298 (*phtV*), with no evidence that it spread across the genome multiple times.

Phylogenetic analysis revealed clear relationships among the Pht cluster sequences. Sequences of *P. syringae* were grouped into 27 haplotypes (that is, DNA sequences that were 100% identical) categorized into three clades: TOX1, TOX2, and TOX3 (Fig. 2). Clades showed divergence, with the prolific TOX1 containing 25 closely related haplotypes (C – X), including all Pph and Psa strains. TOX2 and TOX3 each included a single divergent haplotype (B and A, respectively). Haplotype A was found in Pca ICMP7496, isolated in 1981 from *Carica papaya* in Brazil. Haplotype B was found in Psy strains CFBP3388 and 3023, isolated in 1992 from *Vicia sativa* in France and in 1950 from *Syringa vulgaris* in the UK, respectively (Table S1).

**Figure 2.**
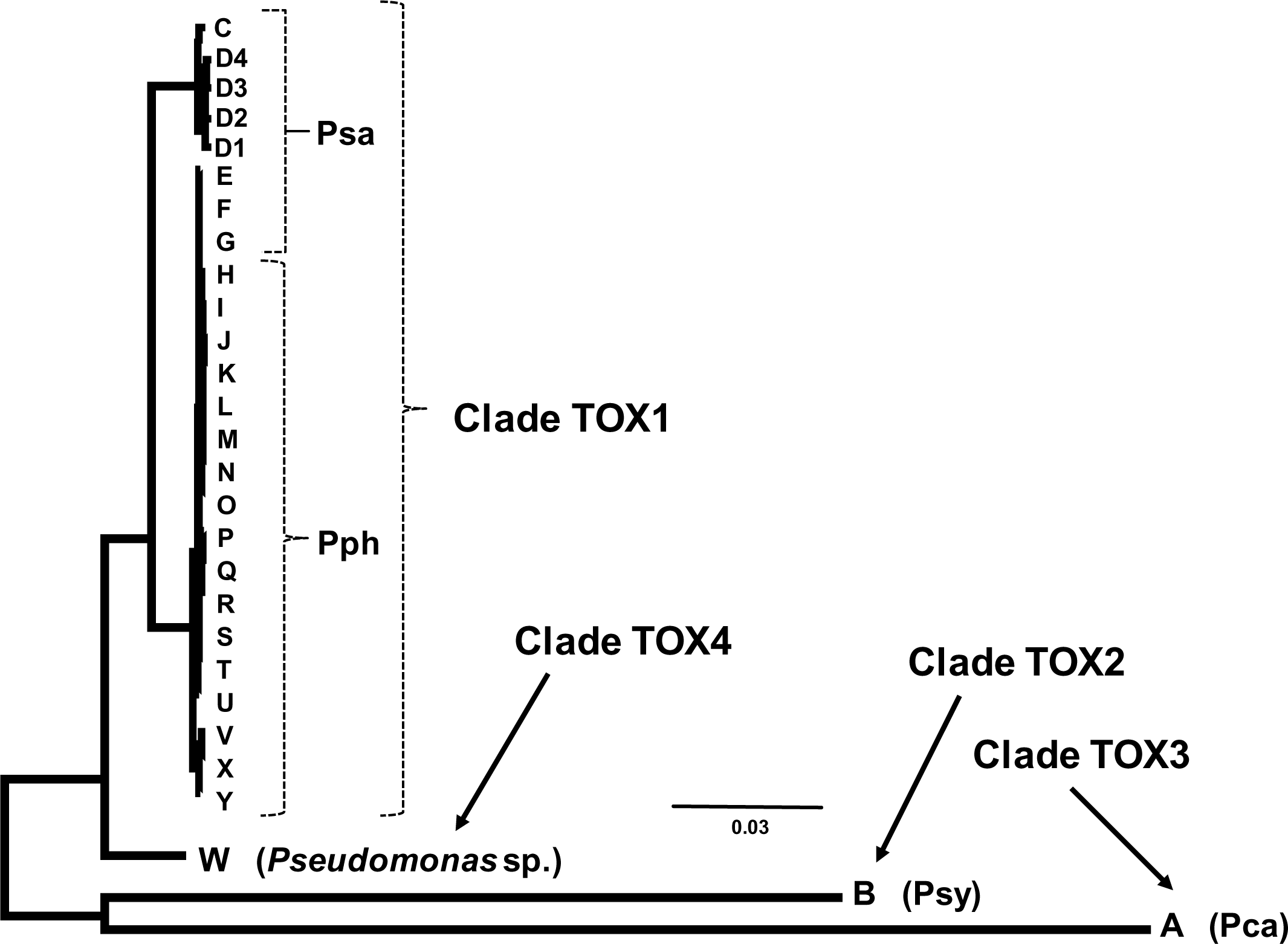
Unrooted Bayesian consensus tree depicting the phylogenetic relationships among the Pht cluster in 102 strains of the *Pseudomonas syringae* complex and *Pseudomonas* sp. JAI115. The dataset (23228 bases, 28 haplotypes) contains DNA sequences of the full Pht cluster, which is located within the chromosomal locus ABC_EAL for all haplotypes, except B and W. Branch lengths are to scale; all branches have 100% posterior probability (not shown). Scale bar indicates expected substitutions per site. The four clades (TOX1 to TOX4) are as depicted. Abbreviations: Psa, *Pseudomonas syringae* pv. actinidiae; Pph, *P. amygdali* pv. phaseolicola; Psy, *P. syringae* pv. syringae; Pca, *P. caricapapayae*.

### The Pht cluster outside *P. syringae*

We also found a fourth TOX clade (haplotype W) in *Pseudomonas* sp. strain JAI115 (Fig. 2), which was obtained from the rhizosphere of the beachgrass *Ammophila breviligulat*a and considered to be a plant growth-promoting bacterium (unpublished; accession no. NZ_JACHCY010000015.1). In a species tree constructed with autoMLST, strain JAI115 clustered together with *P. fluorescens* R124 (ANI 94.4%) and *P. koreensis* (ANI 93.2%) and was distantly related to *P. syringae* (ANI 81.3% with Pph 1448A; 81.9% with Psa ICMP9853; 79.9% with Pca ICMP2855) (data not shown). Thus, TOX4, which consists of haplotype W as its only member, was the sole clade we identified outside *P. syringae*. Interestingly, TOX4 is more closely related to TOX1 than to TOX2 and TOX3 (Fig. 2), with the Pht cluster of strain JAI115 showing 96.9% identity with that of Pph 1448A over 25326 aligned nucleotides.

The Pht cluster in strain JAI115 was found to be part of a large genomic island (Fig. 1), which we identified as a GInt because it conforms to the typical tripartite structure, containing a *ginABCD* operon with three integrases and a small CDS at its 5’ end. Nevertheless, the deduced products of the *ginABCD* operon did not show significant homology with those from Pph strain 1448A in a BLASTP comparison. Additionally, the GInt carrying the Pht cluster in all *P. syringae* strains is specifically integrated at the 5’ end of an ABC transporter. In contrast, in strain JAI115, the corresponding GInt integrated into a different genomic region (Fig. 1).

### Congruence of phylogenies: ABC_EAL locus and Pht cluster in *P. syringae*

We wanted to simultaneously study the evolution of the Pht cluster embedded within the Tox island and that of the genome, as they formed a tandem after the integration event. We thus built a phylogeny using concatenated sequences of two chromosomal genes from the ABC_EAL locus (PSPPH_4293 and PSPPH_4325 homologs) from 330 strains, including toxigenic and nontoxigenic strains. These sequences were condensed into 124 haplotypes. *Pseudomonas* sp. strain JAI115 was not included in this analysis because it is phylogenetically distant from *P. syringae*. The resulting Bayesian phylogenetic tree grouped the 124 haplotypes into six major clades, matching the phylogroups of their parent strains, each with a posterior probability of 100% (Fig. 3). This suggests the ABC_EAL locus evolved alongside the rest of the genome.

**Figure 3.**
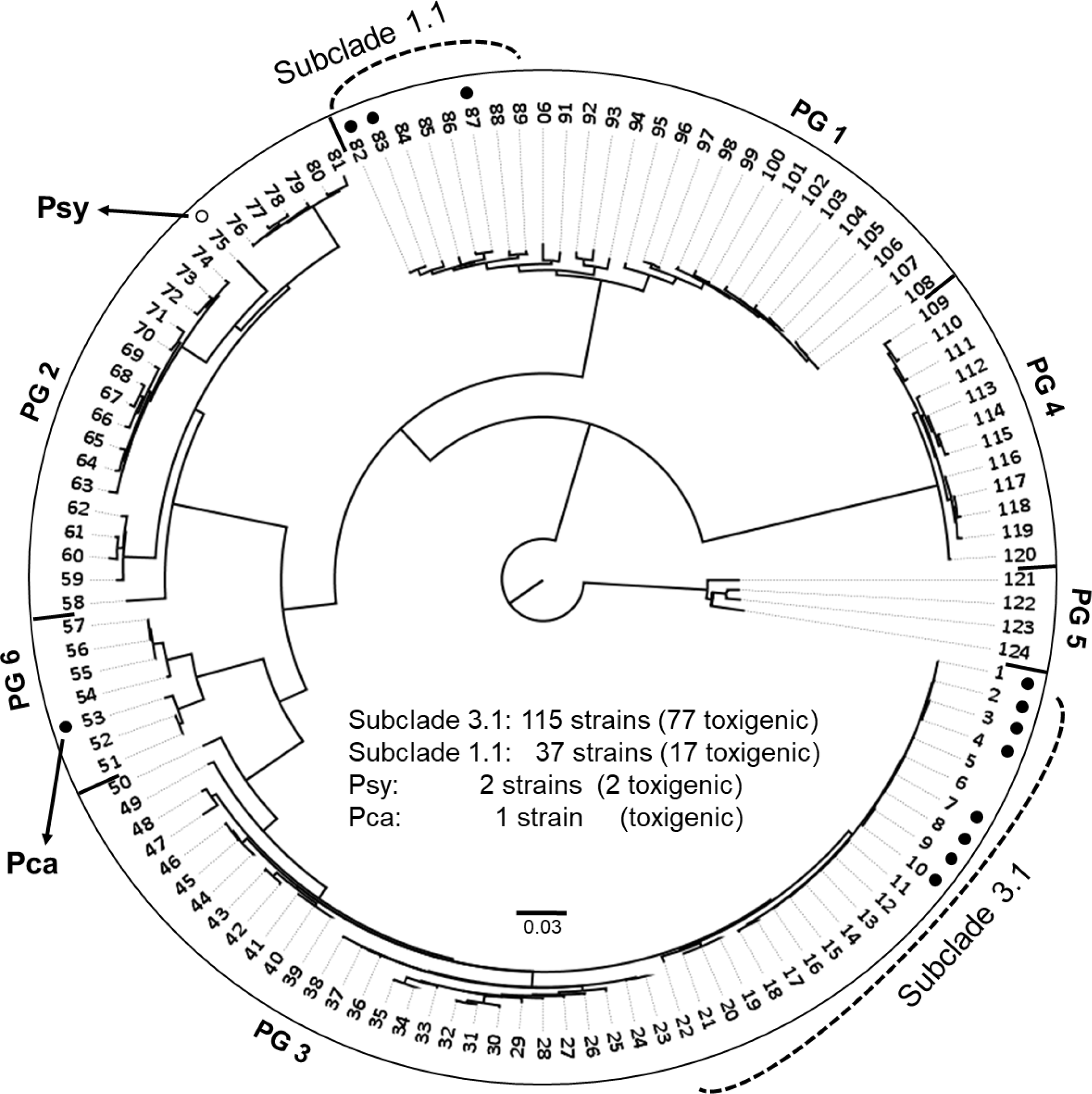
Unrooted Bayesian consensus tree depicting the phylogenetic relationships among the chromosomal locus ABC_EAL in 330 strains of the *Pseudomonas syringae* complex. The dataset (5547 bases, 124 haplotypes) results from merging PSPPH_4293 and PSPPH_4325 (the chromosomal locus ABC_EAL). PSPPH_4293 encodes an ABC transporter; PSPPH_4325 is a pseudogene that otherwise encodes a sensory box/GGDEF domain/EAL domain protein. Haplotypes are color-coded: black circle (Pht cluster within ABC_EAL), open circle (Pht cluster outside ABC_EAL). Branch lengths are to scale; all major branches have 100% posterior probability (not shown). Scale bar indicates expected substitutions per site. PG: Phaseolotoxin-producing phylogroups (Dillon et al., 2019). Abbreviations: Pca, *Pseudomonas caricapapayae*; Psy, *P. syringae* pv. syringae. Strain details in Table S1.

The ABC_EAL locus was occupied by GInts carrying the Pht cluster in three branches: Subclade 3.1 in phylogroup 3 (Pph strains), Subclade 1.1 in phylogroup 1 (Psa strains), and haplotype 53 in phylogroup 6 (Pca ICMP7496) (Figs. 3 and 4). We included both toxigenic and related nontoxigenic strains in Subclades 1.1 and 3.1 for broader phylogenetic context.

**Figure 4.**
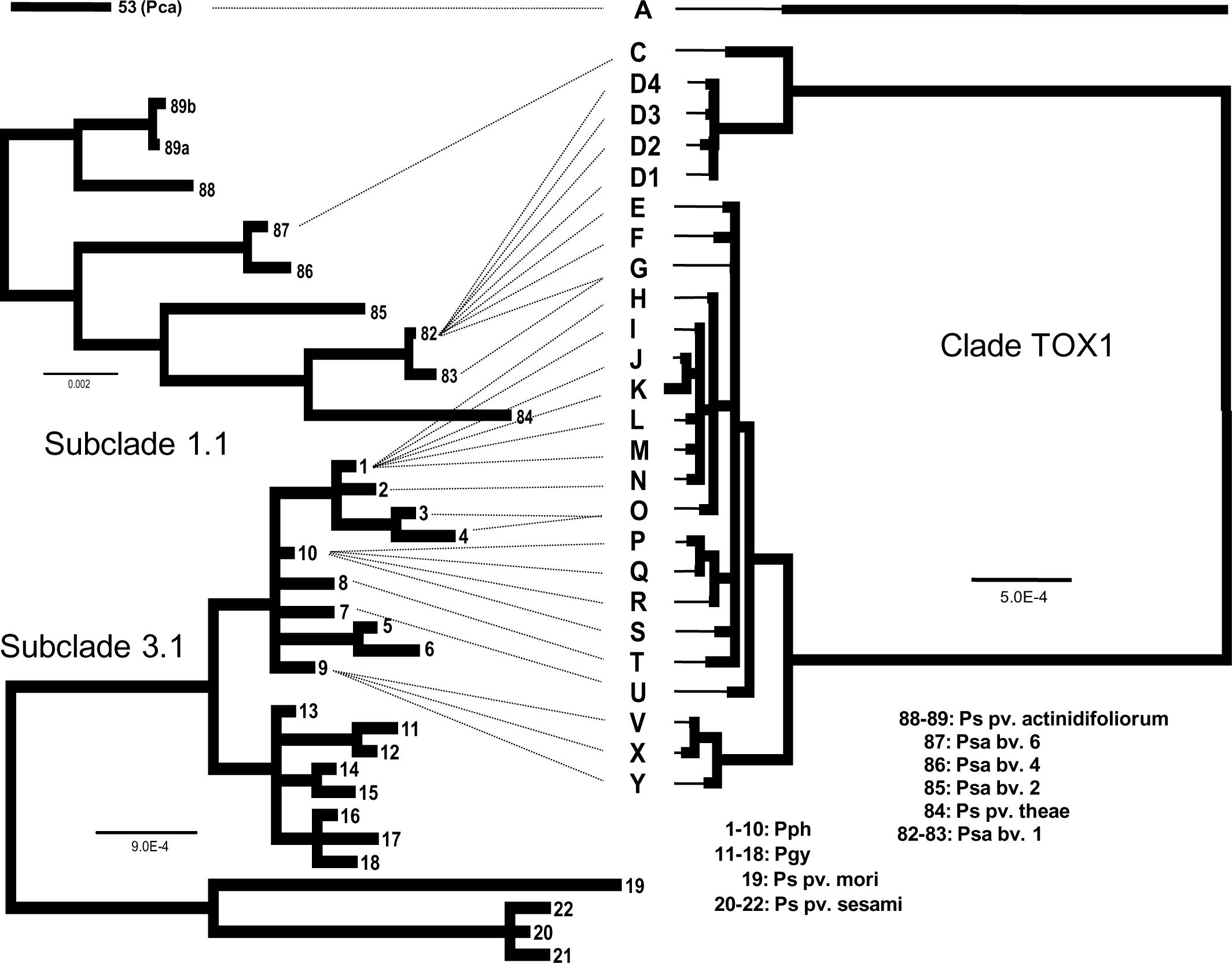
Congruence between the phylogenies of the chromosomal locus ABC_EAL and the Pht cluster. A) Unrooted Bayesian consensus trees of the chromosomal locus ABC_EAL in Subclade 1.1 and Subclade 3.1. The datasets result from merging loci PSPPH_4293 (an ABC transporter) and PSPPH_4325 (a pseudogene that otherwise encodes a sensory box/GGDEF domain/EAL domain protein). These two loci flank the Pht cluster in the two subclades. Subclade 1.1 (top): 5879 bases; 9 haplotypes (45 strains). Subclade 3.1 (bottom): 5880 bases; 22 haplotypes (115 strains). B) Unrooted Bayesian consensus tree of the TOX1 clade. The dataset was 24754 bases, 25 haplotypes. Branch lengths are to scale; all branches have 100% posterior probability (not shown). Scale bar indicates expected substitutions per site. Dotted lines emphasize the congruence between tree topologies. The phylogenetic distant haplotype A (Fig. 3) was not included in the analysis but is shown for reference. Abbreviations: Pca, *Pseudomonas caricapapayae*; Pgy, *P. amygdali* pv. glycinea; Pph, *P. amygdali* pv. phaseolicola; Ps, *P. syringae*; Psa, *P. syringae* pv. actinidiae; Psy, *P. syringae* pv. syringae. pv., pathovar; bv., biovar. Strain details in Table S1.

Subclade 3.1 included 22 haplotypes (Figs. 3 and 4), of which eight corresponded to 77 toxigenic Pph strains (haplotypes 1-4 and 7-10) and eight nontoxigenic Pph strains (haplotypes 5-6). It contained only a small portion of the genetic variation in the ABC_EAL locus (Table 1). Subclade 1.1 had nine haplotypes, including 25 toxigenic Psa strains (Figs. 3 and 4), with higher molecular diversity measures than did Subclade 3.1. This suggests that members of subclade 1.1 underwent diversification to a greater extent (Table 1).

**Table 1.** Measures of molecular diversity.

| Dataset | Component | Size (bp) | n° seq | H | pi | k | S |
| --- | --- | --- | --- | --- | --- | --- | --- |
| **Pht cluster** |  |  |  |  |  |  |  |
| 1 | Toxigenic strain | 23228 | 102 | 28 | 0.028 | 642.4 | 6680 |
| 3 | TOX1 clade* | 24754 | 99 | 25 | 0.00174 | 40.5 | 145 |
| **ABC_EAL** |  |  |  |  |  |  |  |
| 2 | All taxa | 5547 | 330 | 124 | 0.074 | 411.1 | 1842 |
| 4 | Subclade 1.1** | 5879 | 41 | 9 | 0.00630 | 37.0 | 120 |
| 5 | Subclade 3.1 *** | 5880 | 115 | 22 | 0.00134 | 7.8 | 79 |

\*After removing the three most divergent haplotypes (A, B, and W; 4 strains)
\*\*Sequences of the ABC\_EAL locus of *Pseudomonas syringae* pv. actinidiae and closely related taxa
\*\*\*Sequences of the ABC\_EAL locus of *Pseudomonas amygdali* pv. phaseolicola and closely related taxa
H=number of haplotypes; pi=nucleotide diversity; k=average number of nucleotide differences; S=number of variable sites

Haplotype 53 of the ABC_EAL locus originated from one Pca strain (ICMP7496) in phylogroup 6 (Fig. 3), with its Pht cluster belonging to divergent haplotype A (Fig. 2).

### Gene genealogy of the ABC_EAL locus in *P. syringae*

Measures of molecular diversity revealed high polymorphism among Psa strains and related taxa, with a value of pi (average number of nucleotide differences per site) nearly five times greater than that among Pph strains and related taxa (Table 1).

Haplotype networks are similar to phylogenies; however, they offer additional details about the genealogical history of sequences, including ancestral-descendant relationships and haplotype abundance, which may not be evident in phylogenetic analyses. We constructed two networks to visualize the genealogical relationships of the ABC_EAL loci among the Psa and Pph strains (Fig. 5).

**Figure 5.**
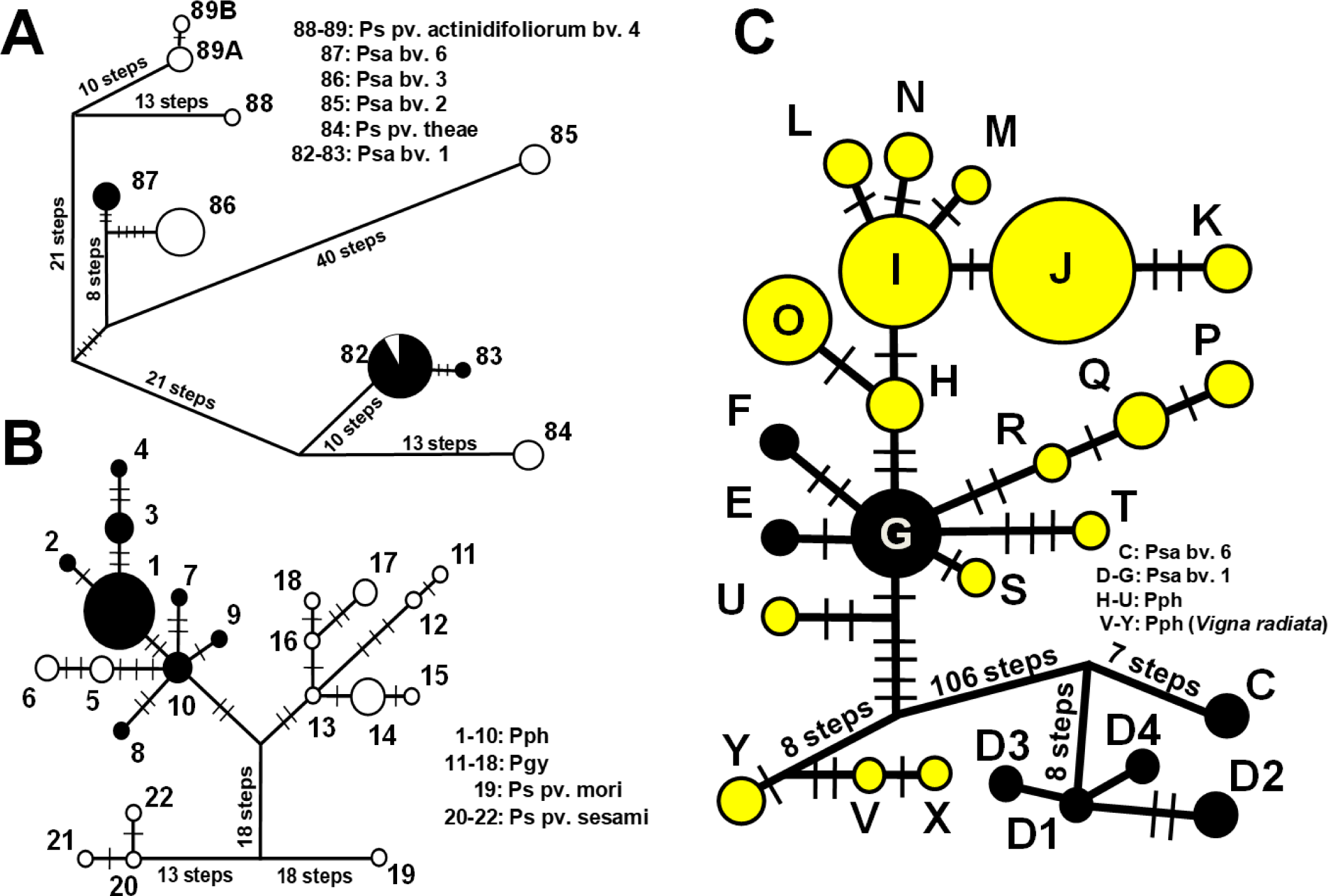
Median-joining network for the chromosomal locus ABC_EAL and the clade TOX1 of the Pht cluster. In each network, a circle represents a given haplotype (coded with numbers according to the phylogenies) (see Fig. 4); circle sizes are proportional to the relative frequencies. Number of mutational steps are indicated with bars when above 5 (unless otherwise indicated). A) Subclade 1.1 in Psa and related pathovars (45 strains, 9 haplotypes, 5879 bases). B) Subclade 1.1 in Pph and related pathovars (115 strains, 22 haplotypes, 5880 bases). Each dataset results from merging PSPPH_4293 and PSPPH_4325. PSPPH_4293 encodes an ABC transporter; PSPPH_4325 is a pseudogene that otherwise encodes a sensory box/GGDEF domain/EAL domain protein. These two regions flank the Pht cluster in toxigenic strains of Psa and Pph. C) Clade TOX1 in Pph (99 strains, 25 haplotypes, 24754 bases). Haplotypes are color-coded: A and B) black circle, Pht cluster within ABC_EAL (toxigenic strains); open circle, the ABC_EAL locus does not contain the Pht cluster (nontoxigenic strains). C) Black circle, Pht cluster of Psa; yellow circles, Pht cluster of Pph. Abbreviations: Pca, *Pseudomonas caricapapayae*; Pgy, *P. amygdali* pv. glycinea; Pph, *P. amygdali* pv. phaseolicola; Ps, *P. syringae*; Psa, *P. syringae* pv. actinidiae; Psy, *P. syringae* pv. syringae. pv., pathovar; bv., biovar. Strain details in Table S1.

The first network, consisting of nine haplotypes from Subclade 1.1, included sequences from Psa, *P. syringae* pv. actinidifoliorum, and *P. syringae* pv. theae, which lack the Pht cluster but share a close relationship with Psa (Fig. 5A). This network revealed high nucleotide diversity, with notable separation between haplotypes, such as the 45 mutational steps between Psa1 (haplotype 82) and Psa6 (haplotype 87).

The second network, comprising 22 haplotypes from Subclade 3.1, included sequences from Pph (toxigenic and nontoxigenic strains) and three nontoxigenic pathovars: *P. syringae* pv. glycinea, mori, and sesami (Fig. 5B). The network showed limited diversity separating the different haplotypes. Thus, the toxigenic Pph strains showed low diversity, with all eight closely related haplotypes descending from the central haplotype 10, which included nine Pph strains (Table S1). Notably, haplotypes 5 and 6 consisted of all the nontoxigenic Pph strains and also descend from haplotype 10, which is specific to toxigenic Pph strains. Haplotype 13 led to a subgroup with eight members from *P. syringae* pv. glycinea and was separated from Pph by 4 mutational steps. Connections to the empty ABC_EAL loci of *P. syringae* pv. sesami and mori required 33 or more mutational steps (Fig. 5B). These three taxa do not contain the Pht cluster.

### Gene genealogy of the clade TOX1 in *P. syringae*

Measures of molecular diversity (e.g., variable sites, average nucleotide differences, and nucleotide diversity) revealed high polymorphism in dataset 1, comprising the entire Pht cluster (Table 1). The derived dataset 3 excluded the three most divergent haplotypes (A, B, and W) and significantly reduced diversity measures by an order of magnitude, indicating a high level of relatedness among TOX1 members.

We constructed a network (Fig. 5C) with Psa (black circles) and Pph strains (yellow circles) to explore the genealogy among TOX1 members. This confirmed the relationships we had uncovered with the phylogenetic reconstruction (Fig. 2) and offered further insights into the genealogical relationships among the members of the TOX1 clade. The genealogy revealed three key findings: 1) Notable differentiation of haplotype C (Psa6) and haplogroup D (Psa1), requiring many mutational steps to connect to the rest of the network. 2) The central role of haplotype G as an ancient ’hub’ from which haplotypes E to T evolved. 3) Independent evolutionary paths of haplotypes V to X (from Pph strains isolated from mung bean, *Vigna radiata*), though less pronounced than those of either C or haplogroup D.

### Congruence of phylogenies: *ginABCD* genes and Pht cluster in *P. syringae*

We evaluated the possible exchange of the Pht cluster among different GInts by examining the congruence of their respective phylogenies.

In all cases, the branches had robust support, and their order agreed in the tree constructed with the concatenated *ginABCD* genes with those in the tree constructed with the 23 concatenated genes of the Pht cluster that they were carrying (Fig. 6). Thus, the divergent position of the *ginABCD* genes of Pca (haplotype I) mirrored the position of haplotype A, the most divergent Pht cluster sequence. Likewise, branching of GInts with haplotypes II and III mirrored the divergent Pht haplotype C and haplogroup D, respectively, from the Psa genomes. Congruence is particularly evident with the GInts from the prolific clade TOX1: GInt haplotypes IV to IX linked, with similar branching, to members of TOX1 (haplotypes E-Y) that were found in the Psa1 and Pph genomes.

**Figure 6.**
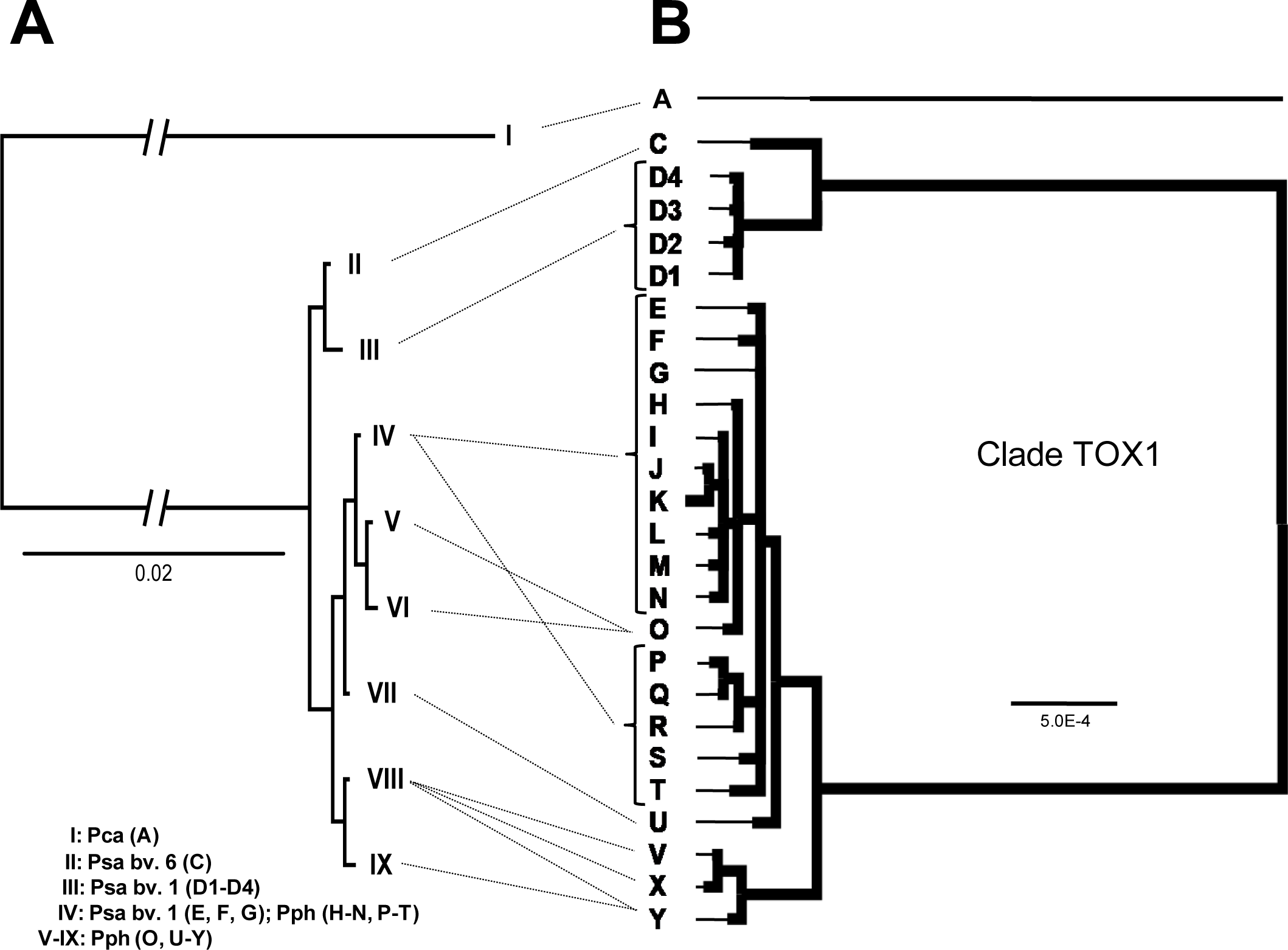
Congruence between the phylogenies of the *ginABCD* genes and the Pht cluster. A) Unrooted Bayesian consensus trees of the *ginABCD* genes. The dataset was 5318 bases long (97 strains; 9 haplotypes) and resulted from merging regions PSPPH_4294 to PSPPH_4297. B) Unrooted Bayesian consensus tree of the TOX1 clade. The dataset was 24754 bases, 25 haplotypes. The phylogenetic distant haplotype A was not included in the analysis but is shown for reference. Branch lengths are to scale; all branches have 100% posterior probability (not shown). Scale bar indicates expected substitutions per site. Dotted lines emphasize the congruence between tree topologies. Abbreviations: Ps, *Pseudomonas syringae*; Pph, *P. amygdali* pv. phaseolicola; Psy, *P. syringae* pv. syringae; Psa, *P. syringae* pv. actinidiae. pv., pathovar; bv., biovar. Strain details in Table S1.

These findings suggest that the *ginABCD* genes and the Pht cluster evolved in tandem over time, indicating a long period of joint evolution during which both gene sets accumulated polymorphisms.

### The *ginABCD* phylogeny within the context of nontoxigenic species in *P. syringae*

The highly divergent GInts found in Pca (TOX2) and in Pph/Psa (TOX1) could either have a distant common ancestor or, alternatively, represent independent acquisitions of the Pht cluster. Therefore, our final phylogenetic reconstruction examined the relationships between the *ginABCD* genes in toxigenic species of *P. syringae* and those in other nontoxigenic strains and species (Fig. S3).

Three key features emerged. First, the *ginABCD* genes carrying the Pht cluster of toxigenic Psa and Pph formed a single, compact clade, indicating low differentiation among members and high differentiation from other clusters in the tree. Second, the highly divergent *ginABCD* genes of Pca (haplotype I) were closely related to those of nontoxigenic *Pseudomonas poae*. Finally, nontoxigenic Pph strains that lost the Pht cluster (holders of haplotypes 5 and 6 of the ABC_EAL locus; Figs. 4 and 5) had *ginABCD* genes distinct from those of toxigenic Pph strains and more closely related to those of *Stutzerimonas zhaodongensis* and other *P. amygdali* strains.

These results indicate that the Pht cluster was independently captured by the GInts present in clades TOX1 and TOX2. Additionally, together with the phylogenetic data of the ABC_EAL locus (Fig. 5), these findings show that the nontoxigenic Pph strains arose by the removal of the GInt carrying the Pht cluster and its replacement with a distantly related GInt carrying a different cargo DNA.

## Discussion

Diverse virulence determinants of plant and animal pathogens, such as those involved in the biosynthesis of secondary metabolites, are inherited horizontally as entire gene clusters. A comparative genomics approach supported by robust phylogenies and network analyses revealed the complex evolutionary history of a 23-gene cluster for the biosynthesis of the phytotoxin phaseolotoxin (the Pht cluster), which serves as a model for understanding the acquisition, evolution, and diversification of genomic islands and virulence factors.

### The Pht cluster is present in five bacterial species and has four configurations

We identified the Pht cluster in five bacterial taxa: *Pseudomonas* sp. JAI115 (related to *P. koreensis*), *P. caricapapayae*, *P. syringae* pv. actinidiae, *P. amygdali* pv. phaseolicola, and *P. syringae* pv. syringae. These clusters likely originated from distinct immediate sources due to their high levels of genetic variation (Fig. 3) and their association with different GInts (Fig. 1). GInts typically transfer DNA among closely related organisms (Smillie et al., 2010; Jackson et al., 2011; Bardaji et al., 2017), suggesting that these clusters were inherited from pseudomonads or related bacteria. Transfers also likely occurred from Pph to Psa1 and from an unidentified source to Psa1 (TOX1 haplotypes D1-D4). Overall, there were at least seven independent acquisitions of this virulence determinant among pseudomonads.

We consistently found a complete Pht cluster with identical gene order and orientations, contrasting with earlier reports of partial presence in *P. syringae* strains (Dillon et al., 2019). This is likely due to our stringent BLAST strategy excluding sequences from the related ’pseudophaseolotoxin region’ (Joardar et al., 2005), which has a rearranged set of 15 homologous genes and is not involved in phaseolotoxin biosynthesis. The 103 Pht cluster sequences identified were grouped into four major clades (TOX1-TOX4) based on phylogeny, GInt type, and insertion site (Figs. 1 and 3).

The Pht cluster in clade TOX1 included 98 strains across 25 closely related haplotypes (C-Y) and is associated with a GInt, termed GInt1, inserted in the chromosomal locus ABC_EAL in Pph and Psa (bv. 1 and bv. 6). The high number of strains in this clade is likely due to sampling bias and sequencing of relevant pathogenic strains, as *P. syringae* often causes clonal epidemics (Sarkar and Guttman, 2004; Arnold et al., 2011; McCann et al., 2017). The rarity of clades TOX2, TOX3, and TOX4 may be due to limited data from natural populations, suggesting undiscovered configurations or clades.

The Pht cluster in clade TOX2 was found in a single Pca strain and was captured by a GInt homologous to GInt1, termed GInt2, also inserted in ABC_EAL. The *ginABCD* genes in TOX2 were more similar to those in non-Pht carrying GInts than to those in TOX1, and the Pht cluster orientation is inverted (Fig. S3).

The Pht cluster in clade TOX3 is in a different chromosomal region, not associated with a mobile genetic element (Fig. 1). It was found in two Psy strains isolated 42 years apart in France and the UK from lilac and a healthy vetch plant (Tourte and Manceau, 1995; Dillon et al., 2019). Their limited presence suggests that they might be environmental strains that do not cause relevant epidemics.

The Pht cluster in clade TOX4 is within a unique DNA region, identified as GInt4, lacking adjacent tRNAs or repeated sequences. The GInt4 structure includes three tyrosine integrases and a small CDS with a helix-turn-helix domain, suggesting its role in integration. While GInt families vary greatly, the GInt of strain JAI115 differs significantly from those of the TOX1 and TOX2 clades, and its insertion site is distinct. This clade marks the third capture of the Pht cluster by a GInt. Additionally, the presence of the Pht cluster in strain JAI115 suggested that it might contribute to environmental survival by outcompeting other organisms in the rhizosphere.

The recurring capture of the Pht cluster into GInt1, GInt2, and GInt4, and the specificity of the MGEs carrying it, is surprising. However, changing the delivery MGE could facilitate the invasion of a wider range of bacteria, as the integration/excision process narrows the host range of GInts (Bardaji et al., 2017). The repeated capture of the cluster is unlikely to be due to general recombination or be mediated by transposable elements, as there are no repeated sequences flanking the cluster. Therefore, it likely involves site-specific recombination mediated by GInt integrases, similar to the mechanism used by integrons (Rowe-Magnus and Mazel, 2002). Nevertheless, identifying conserved sequences mediating this capture is challenging due to the short and variable nature of integrase recognition sequences and recombination sites (Rowe-Magnus and Mazel, 2002; Bardaji et al., 2017).

### The dynamics of Pht cluster acquisition within *P. syringae*

Evidence suggests that the Pht cluster originated from distantly related bacteria and was horizontally transferred into *P. syringae* after pathovar differentiation (Sawada et al., 2002); nevertheless, our study did not focus on the early events of Pht cluster assembly and distribution. Based on previous phylogenetic and structural studies (Sawada et al., 2002; Genka et al., 2006; Murillo et al., 2011; Bardaji et al., 2017; Fujikawa and Sawada, 2019), these events are hypothesized in Fig. 7 (steps 1-6). The phylogeny of the Pht clusters (Fig. 3) suggested that clades TOX2, TOX3, and TOX4 were the first to invade *P. syringae* or other pseudomonads. However, we lack evidence to confirm whether these clusters represent ancient acquisitions that evolved within *P. syringae* or recent transfers of genomic islands. Thus, we focused on the evolution of clade TOX1.

**Figure 7.**
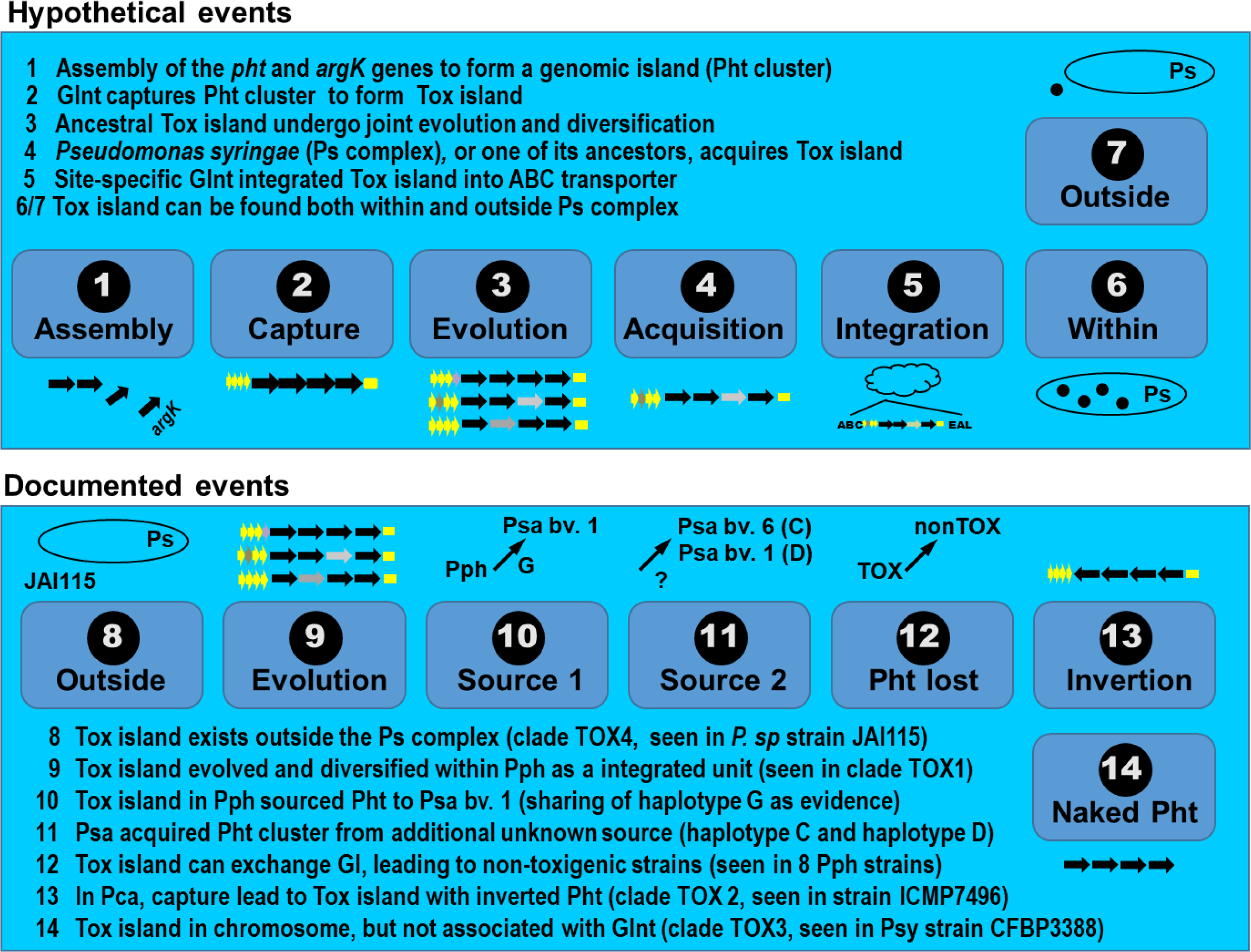
Workflow summarizing hypothetical and documented events outlining the evolutionary history of the phaseolotoxin biosynthesis cluster (Pht cluster).

Previous research identified nearly identical Pht clusters in strains of Pph and Psa (clade TOX1), despite their phylogenetic distance and assignment to different species (Genka et al., 2006; Murillo et al., 2011; Gomila et al., 2017; Sawada and Fujikawa, 2019). To date, it remains unclear whether these clusters were acquired from the same source or transferred between pathovars.

Our phylogenetic analyses (Fig. 4) revealed high identity between the Pht cluster from Pph and most strains of Psa1 (haplotypes G-F). Most Psa1 strains carry an identical Pht cluster (haplotype G), positioned centrally in a network analysis (Fig. 5). This could suggest that haplotype G is ancestral to both pathovars, with Pph possibly acquiring the Tox island from Psa1. However, this hypothesis requires three events: 1) acquisition by ancestral Pph and evolution to haplotypes U-Y, 2) acquisition of haplotype G by Psa1, and 3) transfer of haplotype G to Pph from Psa1. A more parsimonious explanation is that Pph acquired the Tox island and transferred it to Psa1. This is supported by the significant diversification of the Pht cluster in Pph strains and the occurrence of haplotypes U to Y, which do not descend from haplotype G. Thus, we hypothesize that a GInt carrying the Pht cluster (GInt1) invaded an ancestor of Pph early on after pathovar differentiation, as the cluster is absent in *P. amygdali* pv. glycinea (Fig. 2). The acquired GInt1 then diversified within Pph, producing 17 haplotypes of the Pht cluster (Fig. 4). Haplotype G likely existed in Pph but became extinct or unsequenced due to sampling bias, and it was then transferred to an ancestral Psa1, where GInt1 diversified into haplotypes E and F.

Remarkably, we found four Psa1 strains with a highly divergent Pht cluster (haplotypes D1-D4) and four strains without it (Figs. 4 and S4), two of which were previously classified as nontoxigenic by PCR (Sawada et al., 2014). A core genome phylogeny of Psa1 revealed three well-supported clades (haplotypes E-G, haplogroup D, and nontoxigenic) (Fig. S4), but does not clarify the detailed evolutionary history of the Pht cluster in Psa1. All four nontoxigenic strains have a conserved *attB* sequence (5’-CGTATTGAGCC-3’) at the 5’ end of the ABC transporter gene. Since *attB* can be reconstructed after GInt excision (Bardaji et al., 2017), we cannot determine whether these strains are naturally nontoxigenic or if they lost the Tox island. Therefore, the simplest explanation compatible with our data is: an ancestor of Psa1 acquired the Tox island from a Pph strain, forming a clade with haplotypes E-G; later, a nontoxigenic member of this population acquired a different Tox island from an unknown source, generating members of haplogroup D; these lineages coexist with an ancestral nontoxigenic Psa1 lineage.

Finally, Psa6 strains contain a Pht cluster (haplotype C) divergent from the common Psa1 cluster (haplotype G) (Figs. 2 and 5), indicating an additional independent acquisition of the Pht cluster by Psa. However, Psa6 is likely a highly clonal lineage, with a limited distribution (Sawada and Fujikawa, 2019) and few available genomes, preventing a detailed evolutionary analysis.

### Stable configurations postintegration

The abundance of TOX1 clade members highlights relatively straightforward coevolutionary dynamics, showing a tripartite genetic unit that evolved as a single entity: GInt, Pht cluster, and chromosomal locus. The congruence of the phylogenies suggested that this tripartite unit occurred early in evolution for Pph and certain Psa1 strains, evolving as a single entity and accumulating polymorphisms over time.

In Pph, ancestral acquisition led to 17 haplotypes (H to Y). Three haplotypes (V, X, and Y) were divergent and were found in four strains isolated from mung bean, including CFBP3654, which also shows significant core genome divergence from Pph strains from bean (Baltrus et al., 2012; Murillo et al., unpublished). The Pht cluster phylogeny mirrors the ABC_EAL locus phylogeny (Fig. 4), indicating long-term evolution in Pph without recombination or replacement. This stability supports the role of phaseolotoxin in virulence and microbial competition.

Similarly, haplotypes G and D evolved within Psa1 to generate new haplotypes. This genomic stability and the three independent Pht cluster acquisitions support the importance of phaseolotoxin in Psa-plant interactions (Sawada and Fujikawa, 2019).

### Island replacement converts toxigenic strains to nontoxigenic

Nontoxigenic Pph isolates are found in the field, having epidemiological importance in Australia and Spain (Johnson, 1969; Rico et al., 2003), and it is unclear whether they correspond to an ancestral lineage. The ABC_EAL locus in Pph strains forms ten haplotypes, congruent with core-genome phylogenies (Dillon et al., 2019), with eight nontoxigenic strains in haplotypes 5 and 6. Our findings suggest that haplotype 10 is ancestral, from which haplotypes 5 and 6 originated (Figs. 4 and 5B). Nontoxigenic strains contain a homologous but highly divergent GInt in the ABC_EAL locus with different cargo DNAs (Murillo et al., 2011; Bardaji et al., 2017), which is closely related to others in *Stutzerimonas zhaodongensis* and various *P. amygdali* pathovars (Fig. S3). This indicates that nontoxigenic Pph strains may have evolved from toxigenic ancestors that lost the Pht cluster via GInt replacement during evolution and diversification.

### Workflow diagram for the acquisition, evolution, and diversification of the Pht cluster

Our results are summarized in a workflow (Fig. 7) where ‘hypothetical events’ are likely scenarios that stem from our findings, and ‘documented events’ are based on supporting data.

We hypothesized that (1) the 22 *pht* genes for phaseolotoxin biosynthesis and *argK* were assembled from their original donor species. We cannot currently determine the donor’s taxonomic group or whether it represents a single taxon. (2) The Pht cluster, as a genomic island, was captured by a GInt to form the Tox island, which (3) evolved and diversified as a single entity. The mechanism underlying the capture of the Pht cluster by GInts to conform the Tox island remains unknown. (4) Later, a *P. syringae* species, a related congener, or an immediate ancestor then acquired the Tox island. A GInt was integrated site-specifically into the 3’-end of a chromosomal ABC transporter gene in a *P. syringae* species, forming a tripartite genetic unit stably integrated into the genome, either through GInt replacement or by capturing the Tox island as cargo DNA. (6 and 7) Finally, we hypothesize that the Tox island can be found in extant species within or outside the *P. syringae* complex.

Previous analyses hypothesized a relatively straightforward evolutionary scenario for the acquisition of the Tox island by pathovars actinidiae and phaseolicola (Sawada et al., 2002; Genka et al., 2006), two major pathogens of kiwi and bean, respectively. However, detailed phylogenetic and genealogical analyses revealed a very complex evolution of this virulence island in the *P. syringae* complex, which can be described in the following documented steps (Fig. 7):

– **Step 8.** the Tox island existed outside the *P. syringae* complex, which was identified as clade TOX4, in the genome of *Pseudomonas* sp. strain JAI115.
– **Step 9.** The Tox island was acquired by an ancestral Pph strain, and it evolved and diversified as a single unit, together with the genome, integrated into the ABC_EAL locus, as observed for the members of clade TOX1.
– **Step 10.** A Pph strain, which descends from the one that originally acquired the Tox island, transferred this element to a strain of Psa1, generating the lineage of haplotypes E-G of the Pht cluster.
– **Step 11.** Strains of Psa acquired divergent versions of the Tox island from unknown source(s), producing two new lineages:

○ Psa1 containing the D1-D4 haplotypes of the Pht cluster; it is not clear whether this lineage derives from the lineage harboring haplotype G or from a nontoxigenic strain.
○ Psa6 containing haplotype C.
– **Step 12.** We observed the loss of the Tox island in a Pph genetic lineage and its replacement with a divergent GInt that does not carry the Pht cluster. This generated a nontoxigenic lineage that has only been found in Australia and Spain.
– **Step 13.** A different Tox island (clade TOX2) was generated with the capture of the Pht cluster in an inverted orientation by a divergent GInt. This island was either generated in or acquired by *P. caricapapayae* and has only been found in a single strain.
– **Step 14.** The Pht cluster was acquired from an unknown source by *P. syringae* pv. syringae and was integrated into the genome without apparent association with any MGE. It has been found in only two strains.

### The Tox island: a paradigm of genome evolution and bacterial adaptation

GInts (sometimes called Tn6571-family transposons) constitute a highly diverse superfamily present across bacterial phyla (Bardaji et al., 2017; Jiang et al., 2020; Hochhauser et al., 2023). With variable DNA cargo of at least 70 kb, they transport a wide array of genes and gene clusters, often conferring adaptive advantages to microorganisms relevant to human health. Examples include multiresistance to antibiotics in *P. aeruginosa*, a major human pathogen (Moradali et al., 2017), resistance to heavy metals, and genes involved in plant pathogenicity. GInts share structural similarities and limited sequence identity with the Tn*554*-family, which are MGEs associated with multiresistance to antibiotics in *Firmicutes* (Ross et al., 2021); strand-biased circularizing integrative elements, involved in the spread of antimicrobial resistance among *Gammaproteobacteria* (Idola et al., 2023); and recombinase in trio elements, which are widespread in *Bacteria* and are involved in metal resistance (Ricker et al., 2013; Mahbub et al., 2023). Due to their sheer abundance (Audrey et al., 2023; Idola et al., 2023), all these elements constitute a vast and unexplored source of genetic variation in bacteria, whose activity can greatly impact human wellbeing.

It is plausible that all GInts, and similar elements, follow evolutionary paths similar to those of Tox island. Their broad host range and capacity to carry lengthy DNA stretches make them unique models for understanding bacterial evolution and rapid adaptation to challenging environments.

### Conclusions

The phaseolotoxin biosynthesis cluster (the Pht cluster) has been previously found in *P. amygdali* pv. phaseolicola, *P. syringae* pv. actinidiae biovars 1 and 6, and *P. syringae* pv. syringae strain CFBP3388. Here, we report the cluster in strains of two additional species: *P. caricapapayae* ICMP7496, a pathogen of papaya, and *Pseudomonas* sp. JAI115, a plant growth-promoting bacterium isolated from the roots of American beachgrass. The key points that emerged from genomic, phylogenetic and network analyses were that the Pht cluster:

- is present in both plant pathogenic and plant beneficial pseudomonads;
- was found only as a complete unit, including gene *argK*;
- was independently acquired by at least three related but distinct MGEs that are classified as GInts, constituting a virulence island (Tox island);
- has invaded the *P. syringae* complex at least six independent times, originating from unknown sources or from within pseudomonads *P. syringae* members;
- was acquired by *P. syringae* pv. actinidiae at least three independent times, once from *P. amygdali* pv. phaseolicola and two times from unknown sources;
- maintains a stable association with the capturing GInt, with no evidence of recombination among different cluster-carrying GInts;
- once acquired by a bacterium, it tends to be stably maintained, evolving together with the genome by vertical descent, although:

○ the Tox island can be transmitted horizontally to different species of related bacteria;
○ the entire Tox island unit can be replaced by different GInts with dissimilar DNA cargo.
○ the Tox island can be excised from the genome, producing a lineage of nontoxigenic strains.

## Materials and methods

### Bacterial manipulation and molecular biology techniques

Strains were routinely propagated in medium B (KMB) (King et al., 1954) at 28±2 °C, and the production of phaseolotoxin was assayed by an *E. coli* growth inhibition assay, essentially as previously described (Ramírez-Zapata et al., 2020).

Overlapping cosmids containing DNA homologous to the Pht cluster were isolated from a cosmid library of Psy strain CFBP3388 and sequenced by Sanger sequencing of a shotgun library or by primer walking (Macrogen, Seoul, Korea). The resulting fragment (43665 bases) was deposited in the NCBI (accession no. PP449067). Total DNA from CFBP3388 was purified using the Jet Flex Extraction Kit (Genomed; Löhne, Germany), and a 500 bp paired-end library was sequenced using the Illumina platform. Paired reads were qualitatively assessed before de novo assembly with CLC Genomics Workbench (v 10.1.1) software. This Whole Genome Shotgun project has been deposited at DDBJ/ENA/GenBank under the accession JBBAXD000000000. The version described in this paper is version JBBAXD010000000.

### Data acquisition

Genomic data from 268 strains of the *P. syringae* complex were obtained from GenBank. According to a recent phylogenetic reconstruction for this complex (Dillon et al., 2019), these strains were classified into six phylogroups: 1 (69 strains), 2 (36), 3 (96), 4 (21), 5 (10), and 6 (10). We selected these 268 strains based on a previous claim that at least half of the proteins involved in the biosynthetic pathway of phaseolotoxin were present in their individual genomes (Dillon et al., 2019).

Genomic data from 58 other strains of Pph and related pathovars were downloaded from GenBank. Furthermore, 38 additional genomes of Pph and the genome of strain CFBP3388 were included, generated in-house. Finally, we included genomic data for *Pseudomonas* sp. strain JAI115. In total, genomic data from 366 strains were compiled for subsequent analyses.

Local BLASTn searches (E-value < 1e-05) were conducted to identify homologous sequences within the set of 366 strains. For these searches, 33 sequences from Pph strain 1448A (accession no. CP000058) were used as queries. The queries were (1) the complete sequences of the two chromosomal genes (PSPPH_4293 and PSPPH_4325) flanking either side of the integrated GInt that carried the Pht cluster, hereafter referred to as the ABC_EAL locus; (2) nine sequences of the components of the GInt that mediated the integration of the Pht cluster: the four genes PSPPH_4294 to PSPPH_4297 (hereafter referred to as the *ginABCD* genes) and an additional five genes, PSPPH_4320 to PSPPH_4324, located between *argK* and the end of the GInt; and (3) 23 sequences of the components of the Pht cluster extending from gene PSPPH_4319 (*argK*) to gene PSPPH_4298 (*phtV*).

Custom Python scripts were applied to the BLASTn results to retrieve from contigs the full DNA region containing the two chromosomal genes constituting the ABC_EAL locus, along with any integrated sequences that may reside between them. Genomic resources of 36 strains were removed from subsequent analyses because they lacked the target regions within a single contig or had them within two or more contigs for which the ends could not be unambiguously aligned.

Finally, we utilized the DNA sequence of the *ginABCD* genes that are part of the GInt carrying the Pht cluster of *P. caricapapayae* (Pca) strain ICMP7496 as a query during a remote BLASTn search in GenBank. Detailed information about the strains and their descriptions can be found in Table S1.

### Sequence processing and alignments

The retrieved sequences were imported into Sequencher v.4.8 (Gene Codes Corp.) for alignment and editing. Sequencher’s algorithm was employed to conduct multiple-sequence alignments, introducing gaps to accommodate the presence of insertions or deletions (indels). The aligned sequences were trimmed to remove spacer regions between CDSs within the Pht cluster genes. Finally, they were categorized into two sets. One set contained sequences with the full Pht cluster (a total of 103 toxigenic strains). The second set contained the concatenated sequences of the ABC_EAL locus (330 strains), regardless of whether it was empty (224 nontoxigenic strains) or contained integrated sequences (103 toxigenic strains). Sequence data for the ABC_EAL locus of *Pseudomonas* sp. strain JAI115 were not included in these analyses. Sequences of the *ginABCD* genes were aligned using MAFFT online services with default parameters (Katoh et al., 2019). After the alignment and manual trimming processes, several datasets were generated, each of which was meticulously curated to conform to the requirements of the subsequent analysis.

### Phylogenetic analyses

For the first Bayesian analysis, we built dataset 1 (N=28 haplotypes; 23228 bases). This dataset encompassed the entire Pht cluster. After excluding ambiguously aligned segments and indels, the resulting matrix contained 102 sequences, each of which comprised 23228 bases. Subsequently, DnaSP v6 (Rozas et al., 2017) was used to collapse the 102 sequences into 28 haplotypes. Phylogenetic analyses were conducted using Bayesian inference methods implemented in MRBAYES v.3.1.2 (Ronquist and Huelsenbeck, 2003).

Dataset 2 (N=124; 5547 bases) was constructed by concatenating sequences of the chromosomal locus ABC_EAL. Initially comprising 330 sequences and 5547 bases, DnaSP collapsed the sequences into 124 haplotypes.

Building on the results of the two initial analyses, three additional datasets were prepared to investigate the extent to which the phylogeny of the ABC_EAL locus was congruent with the phylogeny of the Pht cluster it harbored.

Dataset 3 (N=25; 24754 bases) contained the entire Pht cluster, but differed from dataset 1 because: (1) it lacked data from the three most highly divergent haplotypes (haplotypes A, B, and W; four strains), and (2) the length of its sequences were larger as the entire region could be aligned unambiguously. DnaSP collapsed the 99 sequences into 25 haplotypes.

Both dataset 4 (N=9; 5879 bases) and dataset 5 (N=22; 5880 bases) contained the concatenated sequences of the chromosomal locus ABC_EAL, but differed from dataset 2. Dataset 4 contained sequences from 45 strains of Psa and the most closely related taxa (Subclade 1.1 in Fig. 3), while dataset 5 harbored sequences from 115 strains of Pph and the most closely related taxa (Subclade 3.1 in Fig. 3). The greater sequence length in those two last clades resulted from the absence of ambiguously aligned regions to be removed after alignment. Subsequently, DnaSP collapsed the sequences of dataset 4 and dataset 5 into nine and 22 haplotypes, respectively.

The concatenated sequences of the *ginABCD* genes formed dataset 6 (N=9; 5318 bases). These sequences were obtained from 99 toxigenic strains within the *P. syringae* complex that had been previously retrieved, which included Pca. In all strains, the *ginABCD* genes were located immediately downstream of the chromosomal ABC transporter gene and upstream of the Pht cluster. DnaSP collapsed the 99 sequences into nine haplotypes.

For the final analysis, we created dataset 7, which contained the 9 haplotypes of the analysis carried out with dataset 6, supplemented with additional sequences of nontoxigenic *P. syringae* strains and other distantly related species. The 34 sequences were retrieved through remote BLAST searches at the NCBI. This analysis was intended to provide insights into the relationships between the *ginABCD* genes present in toxigenic strains and those present in other nontoxigenic strains and species within Ps.

Prior to phylogenetic analysis, each dataset underwent model selection in MRMODELTEST v.2.3 (Nylander, 2004). The Akaike Information Criterion (AIC; Akaike, 1998) identified GTR+I (for datasets 1 and 3) and GTR + I + Γ (for datasets 2-7) as the best-fit models among 24 molecular evolution models. Each analysis comprised two simultaneous runs of three million generations each, with one cold and seven heated chains in each run. The temperature parameter was set to 0.25 (Temp = 0.25), and the branch length prior mean was set to 0.01 (brlenspr = Unconstrained: Exponential (100)). Sampling was performed every 3000 trees, discarding the initial 250 trees as burn-in samples. The average standard deviation of split frequencies at the end of each run was consistently below 0.01, ensuring sufficient posterior sampling. In Tracer 1.5 (Rambaut et al., 2018), Effective Sample Size (ESS) values exceeding 500 for all the statistics indicated reliable sampling. A 50%-majority-rule consensus tree of the two independent runs was obtained, presenting posterior probabilities equal to bipartition frequencies. We used FigTree v1.3.1 (Rambaut, 2009) to visualize the phylogenetic trees.

### Core genome phylogenies

We used the autoMLST web server (https://automlst.ziemertlab.com/analyze) (Alanjary et al., 2019) for the taxonomic placement of *Pseudomonas* sp. JAI115 (accession no. GCF_014200835.1) and for constructing phylogenies of Psa1; we obtained comparable results using the Type Strain Genome Server (https://tygs.dsmz.de/) or Parsnp 1.2 (Treangen et al., 2014), respectively.

### Gene genealogies and molecular diversity estimates

Gene genealogies were inferred from network analyses, having dataset 3, dataset 4, and dataset 5 as independent inputs. We used the median-joining network method (Bandelt et al., 1999), as implemented in NETWORK v10.2.0.0 (Fluxus Technology Ltd.) with default parameters. Measures of molecular diversity (number of haplotypes, H; haplotype diversity, Hd; nucleotide diversity, pi; average number of nucleotide diversity, k; and number of variable sites, S) were estimated in DnaSP v6 (Rozas et al., 2017).

## Availability of data and materials

All genomic data produced in this study have been submitted to the NCBI. Accession numbers for all genomes are included in Table S1.

## Authors’ Contributions

LOO, SA and JM jointly conceived the project. LOO and JM provided the resources. LOO and HVSR designed the pipelines. LOO and JM performed the analyses and interpreted the data. LOO and JM wrote the paper. All authors edited the article and approved the final manuscript.

## Acknowledgments

LOO thanks the Conselho Nacional de Desenvolvimento Científico e Tecnológico (CNPq) for providing a research fellowship (2024-2027) and the Coordenação de Aperfeiçoamento de Pessoal de Nível Superior (CAPES) for providing a Visiting Professor Abroad scholarship under Call No. 01/2023 (Capes/Print-UFV Program for Visiting Professors Abroad 2023 - PPG Quotas). We also extend our gratitude to the Universidade Federal de Viçosa and Universidad Pública de Navarra for facilitating this collaborative research project. We are also thankful to Theresa H. Osinga for help with the English language.

## References

Aguilera, S., López-López, K., Nieto, Y., Garcidueñas-Piña, R., Hernández-Guzmán, G., Hernández-Flores, J.L., et al. (2007). Functional characterization of the gene cluster from *Pseudomonas syringae* pv. phaseolicola NPS3121 involved in synthesis of phaseolotoxin. J. Bacteriol. 189, 2834–2843. doi: 10.1128/JB.01845-06

Akaike, H. (1998). "Information Theory and an Extension of the Maximum Likelihood Principle," in Selected Papers of Hirotugu Akaike, eds. E. Parzen, K. Tanabe & G. Kitagawa. (New York, NY: Springer New York), 199–213.

Alanjary, M., Steinke, K., and Ziemert, N. (2019). AutoMLST: an automated web server for generating multi-locus species trees highlighting natural product potential. Nucleic Acids Res. 47, W276–W282. doi: 10.1093/nar/gkz282

Arndt, H., Henning, C., Völksch, B., and Fritsche, W. (1989). Beziehungen zwischen virulenz und phaseolotoxinbildungsvermögen bei verschiedenen *Pseudomonas syringae* pv. *phaseolicola*-stämmen. Arch. Phytopathol. Pflanzenschutz 25, 347–357. doi: 10.1080/03235408909438888

Arnold, D.L., Lovell, H.C., Jackson, R.W., and Mansfield, J.W. (2011). *Pseudomonas syringae* pv. *phaseolicola*: from ’has bean’ to supermodel. Mol. Plant Pathol. 12, 617–627. doi: 10.1111/j.1364-3703.2010.00697.x

Audrey, B., Cellier, N., White, F., Jacques, P.-É., and Burrus, V. (2023). A systematic approach to classify and characterize genomic islands driven by conjugative mobility using protein signatures. Nucleic Acids Res. 51, 8402–8412. doi: 10.1093/nar/gkad644

Baltrus, D.A., Nishimura, M.T., Dougherty, K.M., Biswas, S., Mukhtar, M.S., Vicente, J., et al. (2012). The molecular basis of host specialization in bean pathovars of *Pseudomonas syringae*. Mol. Plant-Microbe Interact. 25, 877–888. doi: 10.1094/MPMI-08-11-0218

Bandelt, H.J., Forster, P., and Röhl, A. (1999). Median-joining networks for inferring intraspecific phylogenies. Mol. Biol. Evol. 16, 37–48. doi: 10.1093/oxfordjournals.molbev.a026036

Bardaji, L., Echeverría, M., Rodríguez-Palenzuela, P., Martínez-García, P.M., and Murillo, J. (2017). Four genes essential for recombination define GInts, a new type of mobile genomic island widespread in bacteria. Sci. Rep. 7, 46254. doi: 10.1038/srep46254

Bellanger, X., Payot, S., Leblond-Bourget, N., and Guédon, G. (2014). Conjugative and mobilizable genomic islands in bacteria: evolution and diversity. FEMS Microbiol. Rev. 38, 720–760. doi: 10.1111/1574-6976.12058

Bender, C.L., Alarcón-Chaidez, F., and Gross, D.C. (1999). *Pseudomonas syringae* phytotoxins: mode of action, regulation, and biosynthesis by peptide and polyketide synthetases. Microbiol. Mol. Biol. Rev. 63, 266–292. doi: 10.1128%2Fmmbr.63.2.266-292.1999

Berge, O., Monteil, C.L., Bartoli, C., Chandeysson, C., Guilbaud, C., Sands, D.C., et al. (2014). A user’s guide to a data base of the diversity of *Pseudomonas syringae* and its application to classifying strains in this phylogenetic complex. PLoS ONE 9, e105547. doi: 10.1371/journal.pone.0105547

Darmon, E., and Leach, D.R.F. (2014). Bacterial genome instability. Microbiol. Mol. Biol. Rev. 78, 1–39. doi: 10.1128/mmbr.00035-13

Dillon, M.M., Thakur, S., Almeida, R.N.D., Wang, P.W., Weir, B.S., and Guttman, D.S. (2019). Recombination of ecologically and evolutionarily significant loci maintains genetic cohesion in the *Pseudomonas syringae* species complex. Genome Biol. 20, 3. doi: 10.1186/s13059-018-1606-y

Fujikawa, T., and Sawada, H. (2019). Genome analysis of *Pseudomonas syringae* pv. *actinidiae* biovar 6, which produces the phytotoxins, phaseolotoxin and coronatine. Sci. Rep. 9, 3836. doi: 10.1038/s41598-019-40754-9

Genka, H., Baba, T., Tsuda, M., Kanaya, S., Mori, H., Yoshida, T., et al. (2006). Comparative analysis of *argK*-*tox* clusters and their flanking regions in phaseolotoxin-producing *Pseudomonas syringae p*athovars. J. Mol. Evol. 63, 401–414. doi: 10.1007/s00239-005-0271-4

Gomila, M., Busquets, A., Mulet, M., García-Valdés, E., and Lalucat, J. (2017). Clarification of taxonomic status within the *Pseudomonas syringae* species group based on a phylogenomic analysis. Front. Microbiol. 8, 2422. doi: 10.3389/fmicb.2017.02422

Hernández-Morales, A., De la Torre-Zavala, S., Ibarra-Laclette, E., Hernández-Flores, J.L., Jofre-Garfias, A.E., Martínez-Antonio, A., et al. (2009). Transcriptional profile of *Pseudomonas syringae* pv. phaseolicola NPS3121 in response to tissue extracts from a susceptible *Phaseolus vulgaris* L. cultivar. BMC Microbiol. 9, 257. doi: 10.1186/1471-2180-9-257

Hochhauser, D., Millman, A., and Sorek, R. (2023). The defense island repertoire of the *Escherichia coli* pan-genome. PLoS Genet. 19, e1010694. doi: 10.1371/journal.pgen.1010694

Idola, D., Mori, H., Nagata, Y., Nonaka, L., and Yano, H. (2023). Host range of strand-biased circularizing integrative elements: a new class of mobile DNA elements nesting in Gammaproteobacteria. Mobile DNA 14, 7. doi: 10.1186/s13100-023-00295-5

Jackson, R.W., Vinatzer, B., Arnold, D.L., Dorus, S., and Murillo, J. (2011). The influence of the accessory genome on bacterial pathogen evolution. Mob. Genet. Elements 1, 55–65. doi: 10.4161/mge.1.1.16432

Jiang, X., Yin, Z., Yuan, M., Cheng, Q., Hu, L., Xu, Y., et al. (2020). Plasmids of novel incompatibility group Inc_pRBL16_ from *Pseudomonas* species. J. Antimicrob. Chemother. 75, 2093–2100. doi: 10.1093/jac/dkaa143

Joardar, V., Lindeberg, M., Jackson, R.W., Selengut, J., Dodson, R., Brinkac, L.M., et al. (2005). Whole-genome sequence analysis of *Pseudomonas syringae* pv. phaseolicola 1448A reveals divergence among pathovars in genes involved in virulence and transposition. J. Bacteriol. 187, 6488–6498. doi: 10.1128/JB.187.18.6488-6498.2005

Johnson, J.C. (1969). "Halo-less" halo blight of French bean in Queensland. Queensl. J. Agric. Anim. Sci. 26, 293–302.

Katoh, K., Rozewicki, J., and Yamada, K.D. (2019). MAFFT online service: multiple sequence alignment, interactive sequence choice and visualization. Brief. Bioinform. 20, 1160–1166. doi: 10.1093/bib/bbx108

King, E.O., Ward, N.K., and Raney, D.E. (1954). Two simple media for the demonstration of pyocyanin and fluorescin. J. Lab. Clin. Med. 44, 301–307. doi: 10.5555/uri:pii:002221435490222X

Mahbub, K.R., Chénard, C., Batinovic, S., Petrovski, S., Lauro, F.M., Rahman, M.H., et al. (2023). Complex interactions between diverse mobile genetic elements drive the evolution of metal-resistant bacterial genomes. Environ. Microbiol. 25, 3387–3405. doi: 10.1111/1462-2920.16532

Mansfield, J., Genin, S., Magori, S., Citovsky, V., Sriariyanum, M., Ronald, P., et al. (2012). Top 10 plant pathogenic bacteria in molecular plant pathology. Mol. Plant Pathol. 13, 614–629. doi: 10.1111/j.1364-3703.2012.00804.x

McCann, H.C., Li, L., Liu, Y., Li, D., Pan, H., Zhong, C., et al. (2017). Origin and evolution of the kiwifruit canker pandemic. Genome Biol. Evol. 9, 932–944. doi: 10.1093/gbe/evx055

Mitchell, R.E. (1976). Isolation and structure of a chlorosis-inducing toxin of *Pseudomonas phaseolicola*. Phytochemistry 15, 1941–1947. doi: 10.1016/S0031-9422(00)88851-2

Mitchell, R.E. (1978). Halo blight of beans: toxin production by several *Pseudomonas phaseolicola* isolates. Physiol. Plant Pathol. 13, 37–49. doi: 10.1016/0048-4059(78)90073-5

Miyoshi, T., Shimizu, S., and Sawada, H. (2012). Occurrence and distribution of a defective non-phaseolotoxin-producing mutant of *Pseudomonas syringae* pv. *actinidiae* in Ehime Prefecture, Japan. Jpn. J. Phytopathol. 78, 92–103. doi: 10.3186/jjphytopath.78.92

Moradali, M.F., Ghods, S., and Rehm, B.H. (2017). *Pseudomonas aeruginosa l*ifestyle: A paradigm for adaptation, survival, and persistence. Front. Cell. Infect. Microbiol. 7, 39. doi: 10.3389/fcimb.2017.00039

Murillo, J., Bardaji, L., Navarro de la Fuente, L., Führer, M.E., Aguilera, S., and Alvarez-Morales, A. (2011). Variation in conservation of the cluster for biosynthesis of the phytotoxin phaseolotoxin in *Pseudomonas syringae* suggests at least two events of horizontal acquisition. Res. Microbiol. 162, 253–261. doi: 10.1016/j.resmic.2010.10.011

Nylander, J.A.A. (2004). "MrModeltest v2". (Program distributed by the author: Evolutionary Biology Centre, Uppsala University).

Patil, S.S., Hayward, A.C., and Emmons, R. (1974). An ultraviolet-induced non-toxigenic mutant of *Pseudomonas phaseolicola* of altered pathogenicity. Phytopathology 64, 590–595. doi: 10.1094/Phyto-64-590

Rambaut, A. (2009). *FigTree v1.3.1. Institute of Evolutionary Biology, University of Edinburgh, Edinburgh*. [Online]. http://tree.bio.ed.ac.uk/software/figtree/. Available: http://tree.bio.ed.ac.uk/software/figtree/ [Accessed].

Rambaut, A., Drummond, A.J., Xie, D., Baele, G., and Suchard, M.A. (2018). Posterior summarization in Bayesian phylogenetics using Tracer 1.7. Syst. Biol. 67, 901–904. doi: 10.1093/sysbio/syy032

Ramírez-Zapata, D., Ramos, C., Aguilera, S., Bardaji, L., Martínez-Gil, M., and Murillo, J. (2020). Two homologues of the global regulator Csr/Rsm redundantly control phaseolotoxin biosynthesis and virulence in the plant pathogen *Pseudomonas amygdali* pv. phaseolicola 1448A. Microorganisms 8, 1536. doi: 10.3390/microorganisms8101536

Ricker, N., Qian, H., and Fulthorpe, R.R. (2013). Phylogeny and organization of recombinase in trio (RIT) elements. Plasmid 70, 226–239. doi: 10.1016/j.plasmid.2013.04.003

Rico, A., López, R., Asensio, C., Aizpún, M., Asensio-S.-Manzanera, C., and Murillo, J. (2003). Nontoxigenic strains of *P. syringae* pv. *phaseolicola* are a main cause of halo blight of beans in Spain and escape current detection methods. Phytopathology 93, 1553–1559. doi: 10.1094/PHYTO.2003.93.12.1553

Ronquist, F., and Huelsenbeck, J.P. (2003). MrBayes 3: Bayesian phylogenetic inference under mixed models. Bioinformatics 19, 1572–1574. doi: 10.1093/bioinformatics/btg180

Ross, K., Varani, A.M., Snesrud, E., Huang, H., Alvarenga, D.O., Zhang, J., et al. (2021). TnCentral: a prokaryotic transposable element database and web portal for transposon analysis. mBio 12, 10.1128/mbio.02060-02021. doi: 10.1128/mbio.02060-21

Rowe-Magnus, D.A., and Mazel, D. (2002). The role of integrons in antibiotic resistance gene capture. Int. J. Med. Microbiol. 292, 115–125. doi: 10.1078/1438-4221-00197

Rozas, J., Ferrer-Mata, A., Sánchez-DelBarrio, J.C., Guirao-Rico, S., Librado, P., Ramos-Onsins, S.E., et al. (2017). DnaSP 6: DNA sequence polymorphism snalysis of large data sets. Mol. Biol. Evol. 34, 3299–3302. doi: 10.1093/molbev/msx248

Sarkar, S.F., and Guttman, D.S. (2004). Evolution of the core genome of *Pseudomonas syringae*, a highly clonal, endemic plant pathogen. Appl. Environ. Microbiol. 70, 1999–2012.

Sawada, H., and Fujikawa, T. (2019). Genetic diversity of *Pseudomonas syringae* pv. *actinidiae*, pathogen of kiwifruit bacterial canker. Plant Pathol. 68, 1235–1248. doi: 10.1111/ppa.13040

Sawada, H., Kanaya, S., Tsuda, M., Suzuki, F., Azegami, K., and Saitou, N. (2002). A phylogenomic study of the OCTase genes in *Pseudomonas syringae* pathovars: the horizontal transfer of the *argK*–*tox* cluster and the evolutionary history of OCTase genes on their genomes. J. Mol. Evol. 54, 437–457. doi: 10.1007/s00239-001-0032-y

Sawada, H., Miyoshi, T., and Ide, Y. (2014). Novel MLSA group (Psa5) of *Pseudomonas syringae* pv. *actinidiae* causing bacterial canker of kiwifruit (*Actinidia chinensis*) in Japan. Jpn. J. Phytopathol. 80, 171–184. doi: 10.3186/jjphytopath.80.171

Smillie, C., Garcillán-Barcia, M.P., Francia, M.V., Rocha, E.P.C., and de la Cruz, F. (2010). Mobility of plasmids. Microbiol. Mol. Biol. Rev. 74, 434–452. doi: 10.1128/mmbr.00020-10

Tamura, K., Imamura, M., Yoneyama, K., Kohno, Y., Takikawa, Y., Yamaguchi, I., et al. (2002). Role of phaseolotoxin production by *Pseudomonas syringae* pv. *actinidiae* in the formation of halo lesions of kiwifruit canker disease. Physiol. Mol. Plant Pathol. 60, 207–214. doi: 10.1006/pmpp.2002.0405

Tourte, C., and Manceau, C. (1995). A strain of *Pseudomonas syringae* which does not belong to pathovar *phaseolicola* produces phaseolotoxin. Eur. J. Plant Pathol. 101, 483–490. doi: 10.1007/BF01874471

Treangen, T.J., Ondov, B.D., Koren, S., and Phillippy, A.M. (2014). The Harvest suite for rapid core-genome alignment and visualization of thousands of intraspecific microbial genomes. Genome Biol. 15, 524. doi: 10.1186/s13059-014-0524-x

Uddin, S., Bae, D., Cha, J.Y., Ahn, G., Kim, W.Y., and Kim, M.G. (2022). Coronatine induces stomatal reopening by inhibiting hormone signaling pathways. J. Plant Biol. 65, 403–411. doi: 10.1007/s12374-022-09362-5

