## Supplementary Figures for "Acquisition, Evolution, and Diversification of Genomic Islands: A Case Study of a Virulence Gene Cluster in *Pseudomonas syringae*"

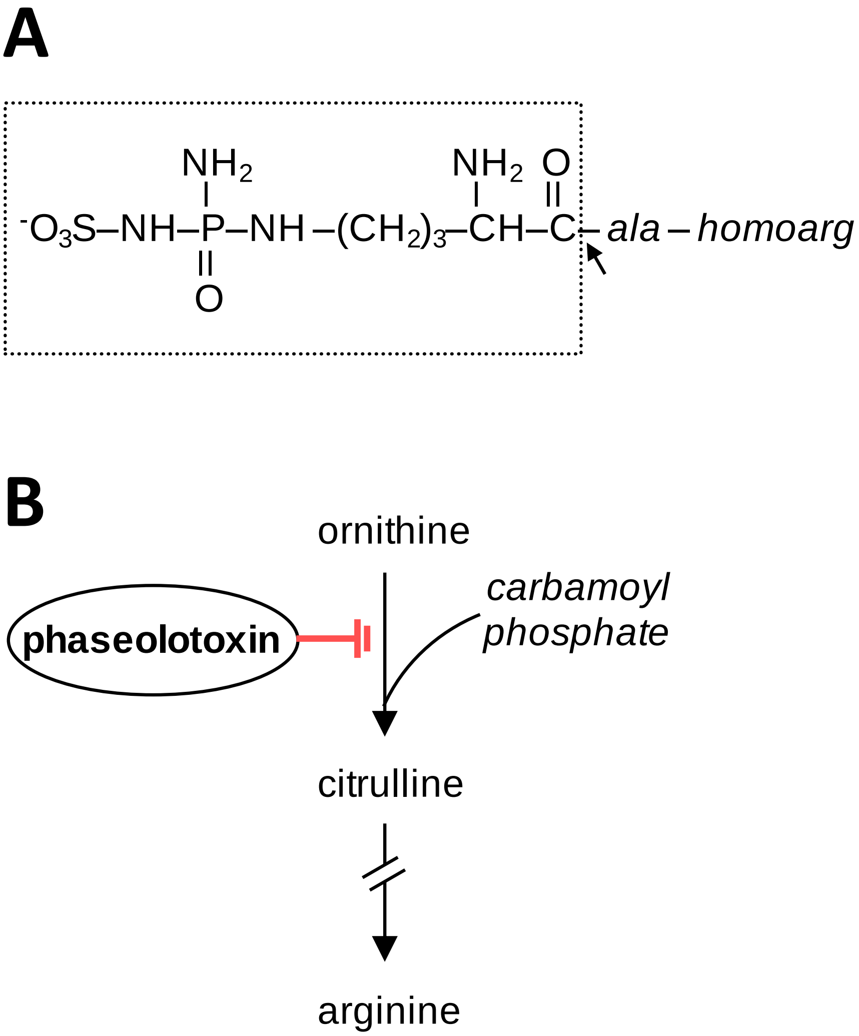


**Figure S1. Structure and activity of phaseolotoxin.** A) Chemical structure of phaseolotoxin. The arrow indicates the point of hydrolysis by peptidases, which liberate the predominant active form of the toxin, octicidin (boxed); homoarg, homoarginine. B) Phaseolotoxin inhibits the enzyme ornithine carbamoyl transferase, a critical enzyme in the arginine biosynthesis pathway that catalyzes the production of L-citrulline from ornithine and carbamoyl phosphate.

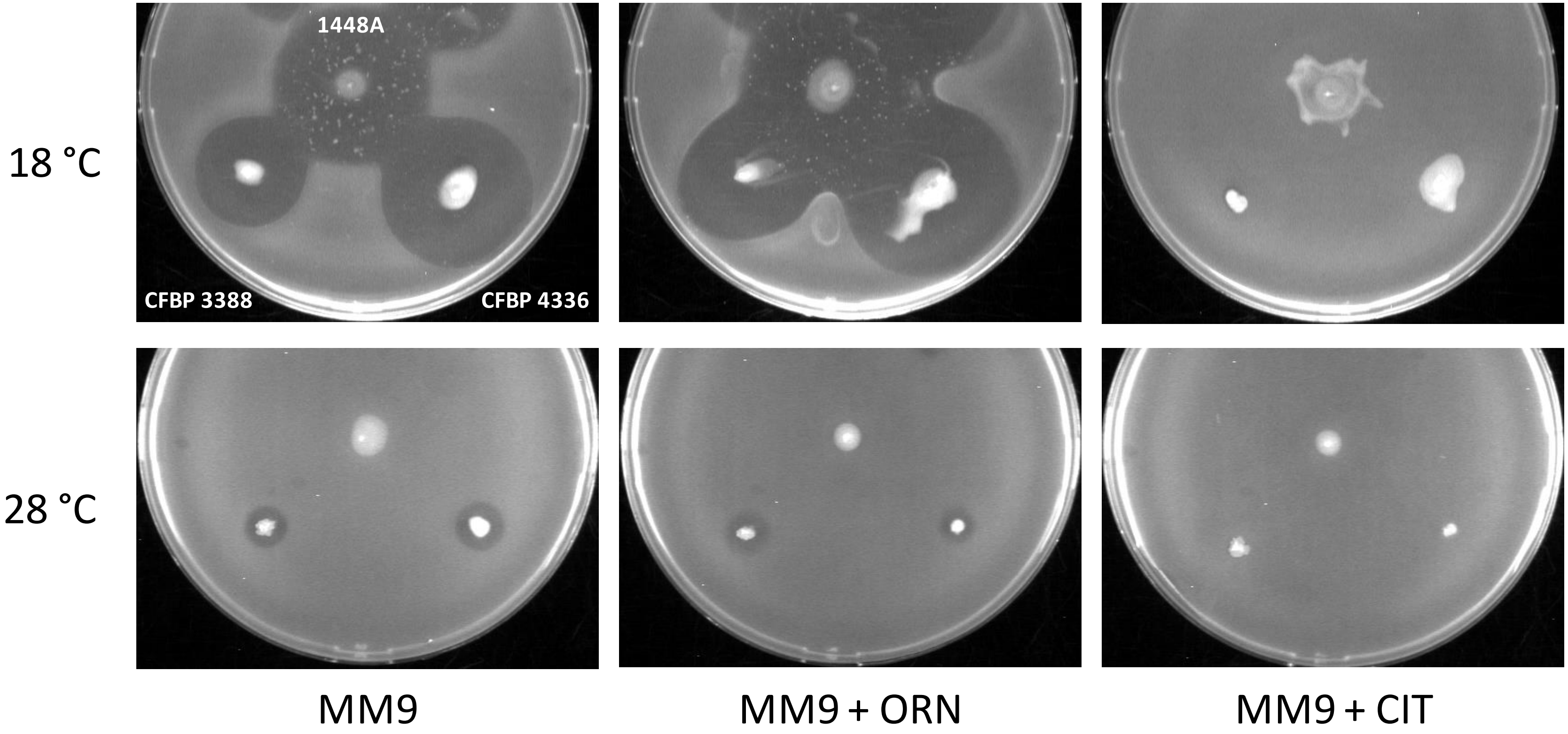


**Figure S2. Production of phaseolotoxin using an *E. coli* growth inhibition assay.** *E. coli* S17-1 λpir was layered onto MM9 using soft agar, which was inoculated with three strains: *P. amygdali* pv. phaseolicola 1448A, *P. syringae* pv. syringae CFBP3388, and *P. caricapapayae* CFBP4336. The plates were incubated at either 18 or 28 ºC. Inhibition of ornithine carbamoyltransferase by phaseolotoxin is specifically evidenced as halos of growth inhibition of *E. coli* on MM9 plates with or without ornithine (ORN), with the reversion of halos on MM9 plates supplemented with citrulline. The experiment was repeated four times with consistent results.

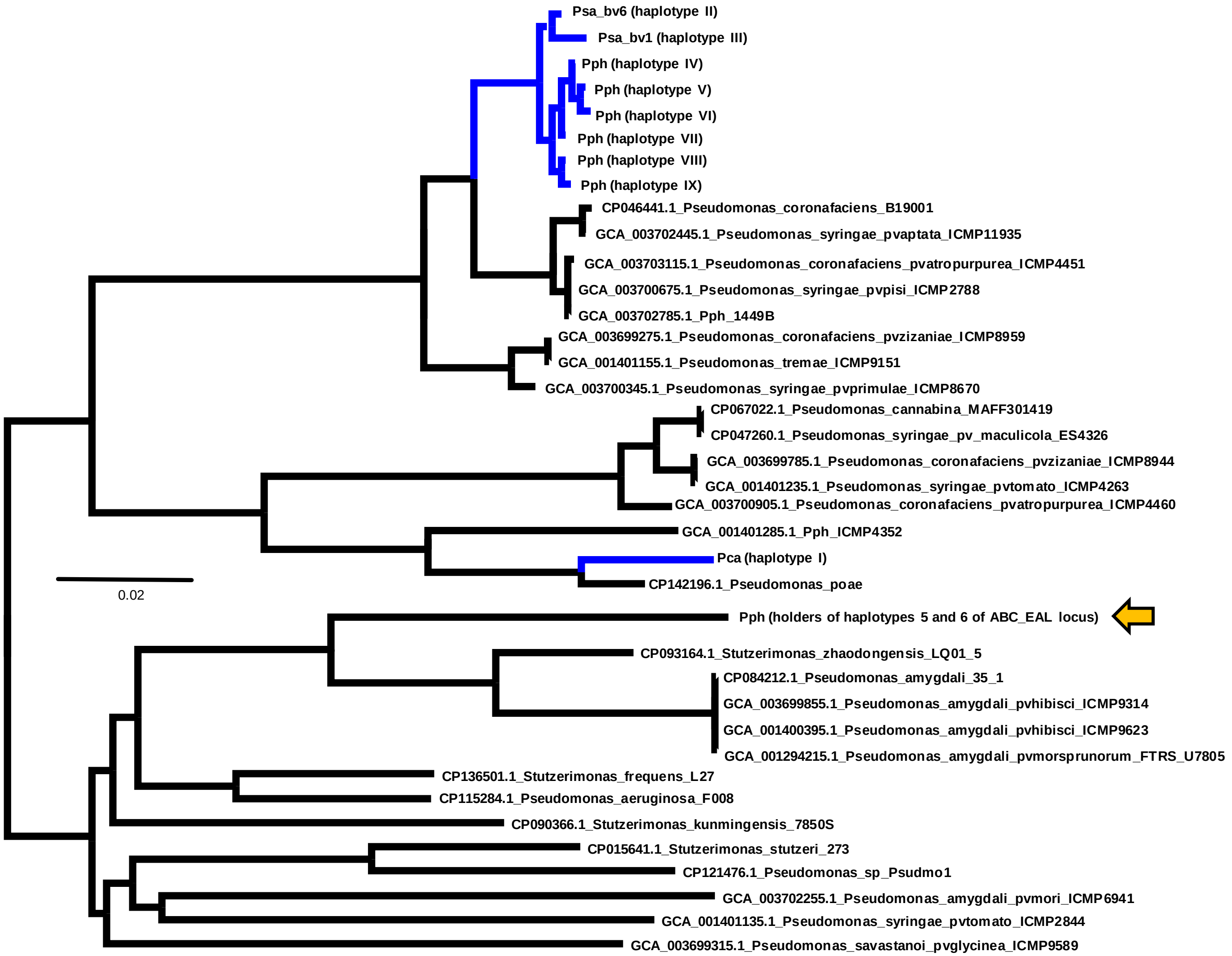


**Figure S3. Unrooted Bayesian consensus tree depicting the phylogenetic relationships among the *ginABCD* genes of strains of the *Pseudomonas syringae* complex and closely related species.** The dataset was 5534 bases long (43 sequences) and resulted from merging PSPPH_4294 to PSPPH_4297. Branch lengths are to scale; all branches have 100% posterior probability (not shown). Scale bar indicates expected substitutions per site. Abbreviations: Ps, *Pseudomonas syringae*; Pph, *P. amygdali* pv. phaseolicola; Psy, *P. syringae* pv. syringae; Psa, *P. syringae* pv. actinidiae. pv., pathovar; bv., biovar. Strain details in Table S1.

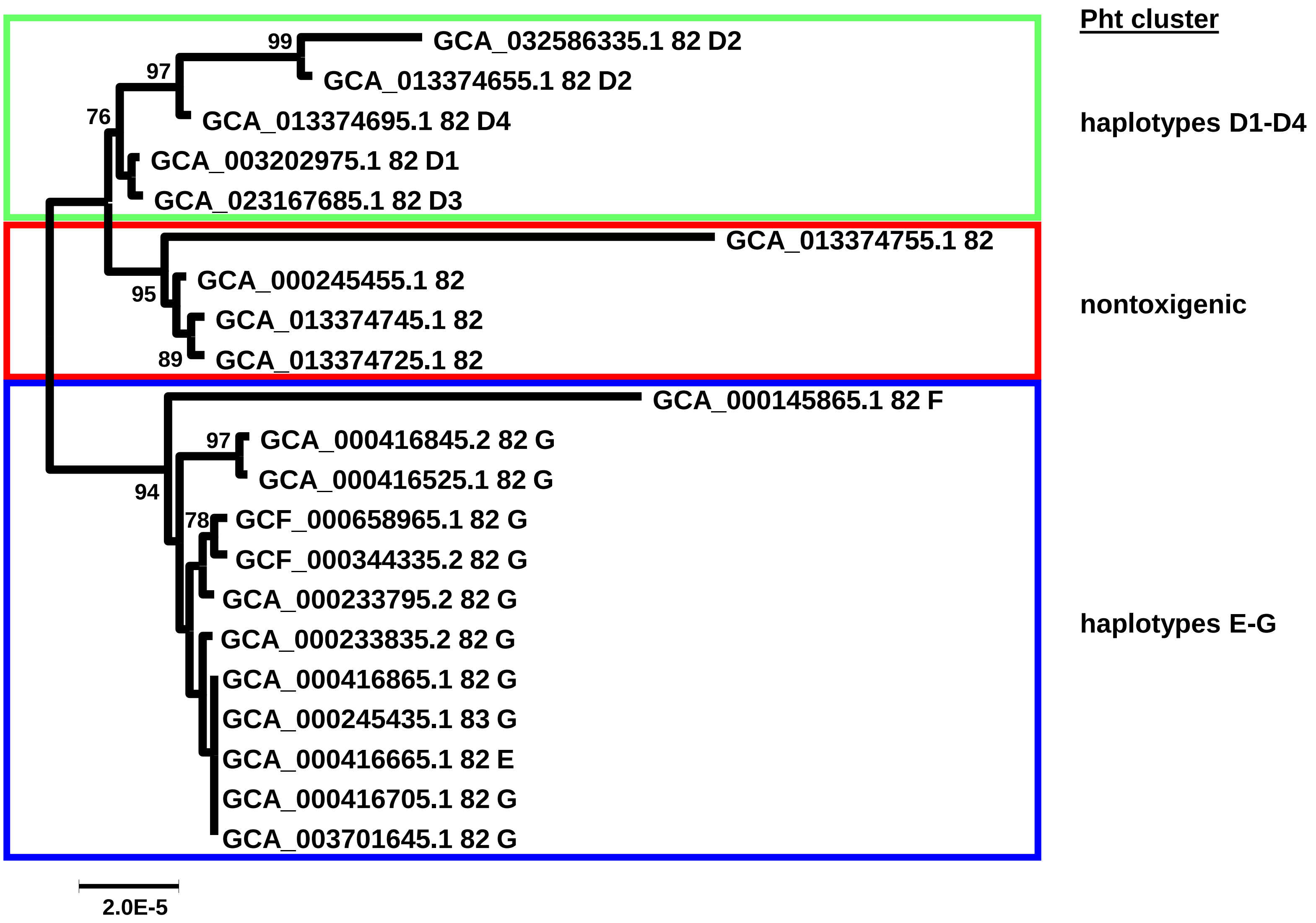


**Figure S4. Maximum likelihood tree of an MLSA analysis of *P. syringae* pv. actinidiae biovar 1 strains.** The analysis was carried out in the autoMLST server (https://automlst.ziemertlab.com/) with default options and used 100 concatenated core genes. Numbers in nodes are percentages of ultrafast bootstrap replicates (shown when above 70). The tree is rooted with closely related strains of *P. syringae sensu lato*, which are not shown for clarity. Following the assembly accession number for each genome are the haplotypes for the ABC-EAL locus and for the Pht cluster. The scale indicates number of changes per nucleotide.
